# CRISPR interference-based platform for multimodal genetic screens in human iPSC-derived neurons

**DOI:** 10.1101/513309

**Authors:** Ruilin Tian, Mariam A. Gachechiladze, Connor H. Ludwig, Matthew T. Laurie, Jason Y. Hong, Diane Nathaniel, Anika V. Prabhu, Michael S. Fernandopulle, Rajan Patel, Mehrnoosh Abshari, Michael E. Ward, Martin Kampmann

## Abstract

CRISPR/Cas9-based functional genomics have transformed our ability to elucidate mammalian cell biology. However, most previous CRISPR-based screens were conducted in cancer cell lines, rather than healthy, differentiated cells. Here, we describe a CRISPR interference (CRISPRi)-based platform for genetic screens in human neurons derived from induced pluripotent stem cells (iPSCs). We demonstrate robust and durable knockdown of endogenous genes in such neurons, and present results from three complementary genetic screens. First, a survival-based screen revealed neuron-specific essential genes and genes that improved neuronal survival upon knockdown. Second, a screen with a single-cell transcriptomic readout uncovered several examples of genes whose knockdown had strikingly cell-type specific consequences. Third, a longitudinal imaging screen detected distinct consequences of gene knockdown on neuronal morphology. Our results highlight the power of unbiased genetic screens in iPSC-derived differentiated cell types and provide a platform for systematic interrogation of normal and disease states of neurons.

## INTRODUCTION

While DNA sequencing has provided us with an inventory of human genes, and RNA sequencing is revealing when and where these genes are expressed, the next challenge is to systematically understand the function of human genes in different cell types. A powerful approach to functionally annotate the human genome is genetic screening in cultured cells. The robustness of such screens has improved substantially through the recent introduction of CRISPR/Cas9-based approaches. Cas9 nuclease can be targeted by single guide RNAs (sgRNAs) to introduce DNA breaks in coding regions of genes, which are subsequently repaired by non-homologous end-joining pathways. This process frequently causes short deletions or insertions that disrupt gene function. This CRISPR nuclease (CRISPRn) strategy has enabled genetic screens through the use of pooled sgRNA libraries targeting large numbers of genes (Koike-Yusa et al., 2014; Shalem et al., 2014; Wang et al., 2014; Zhou et al., 2014). We previously developed an alternative platform for loss-of-function screens in mammalian cells based on CRISPR interference (CRISPRi) (Gilbert et al., 2014). In CRISPRi screens, sgRNAs target catalytically dead Cas9 (dCas9) fused to a KRAB transcriptional repression domain to transcription start sites in the genome, thereby inhibiting gene transcription. CRISPRn and CRISPRi screening platforms each have their advantages for specific applications (Kampmann, 2018; Rosenbluh et al., 2017), but generally yield similar results (Horlbeck et al., 2016). Most previous CRISPR-based screens were implemented in cancer cell lines or stem cells rather than healthy differentiated human cells, thereby limiting potential insights into cell type-specific roles of human genes.

Here, we present a CRISPRi-based platform for genetic screens in human induced pluripotent stem cell (iPSC)-derived neurons. To our knowledge, it is the first description of a large-scale CRISPR-based screening platform in any differentiated, human iPSC-derived cell type. We focused on neurons as our first application, since functional genomic screens in human neurons have the potential to reveal mechanisms of selective vulnerability in neurodegenerative diseases (Kampmann, 2017) and convergent mechanisms of neuropsychiatric disorders (Willsey et al., 2018), thus addressing urgent public health issues. iPSC technology is particularly relevant to the study of human neurons, since primary neurons are difficult to obtain from human donors, and non-expandable due to their post-mitotic nature.

We integrated CRISPRi technology with our previously described i^3^Neuron platform (Fernandopulle et al., 2018; Wang et al., 2017), which yields large quantities of highly homogeneous neurons, a prerequisite for robust population-based screens. We decided to use CRISPRi rather than CRISPRn, since CRISPRn-associated DNA damage is highly toxic to iPSCs and untransformed cells (Haapaniemi et al., 2018; Ihry et al., 2018; Schiroli et al., 2019). Furthermore, CRISPRi perturbs gene function by partial knockdown, rather than knockout, thereby enabling the investigation of the biology of essential genes. While large-scale genetic screens in mouse primary neurons have previously been implemented using RNA interference (RNAi) technology (Nieland et al., 2014; Sharma et al., 2013), CRISPRi represents an important advance over RNAi, since it lacks the pervasive off-target effects (Gilbert et al., 2014) inherent to RNAi-based screening approaches (Adamson et al., 2012; Jackson et al., 2003; Kaelin, 2012).

We demonstrate the versatility of our approach in three complementary genetic screens, based on neuronal survival, single-cell RNA sequencing (scRNA-Seq), and neuronal morphology. These screens revealed striking examples of cell-type specific gene functions and identified new genetic modifiers of neuronal biology. Our results provide a strategy for systematic dissection of normal and disease states of neurons, and highlight the potential of interrogating human cell biology and gene function in iPSC-derived differentiated cell types.

## RESULTS

### Robust CRISPR interference in human iPSC-derived neurons

As a first step towards a high-throughput screening platform in neurons, we developed a scalable CRISPRi-based strategy for robust knockdown of endogenous genes in homogeneous populations of human iPSC-derived neurons. We built on our previously described i^3^Neuron (i^3^N) platform, which enables large-scale production of iPSC-derived glutamatergic neurons. Central to this platform is an iPSC line with an inducible Neurogenin 2 (Ngn2) expression cassette (Zhang et al., 2013) in the AAVS1 safe-harbor locus (Fernandopulle et al., 2018; Wang et al., 2017). To enable stable CRISPRi in iPSC-derived neurons, we generated a plasmid (pC13N-dCas9-BFP-KRAB) to insert an expression cassette for CAG promoter-driven dCas9-BFP-KRAB into the CLYBL safe harbor locus, which enables robust transgene expression throughout neuronal differentiation at higher levels than the AAVS1 locus (Cerbini et al., 2015) (Fig. 1A). We then integrated this cassette into our i^3^N iPSC line, and called the resulting monoclonal line CRISPRi-i^3^N iPSCs. A normal karyotype was confirmed for CRISPRi-i^3^N iPSCs (Fig. S1A).

**Fig. 1.**
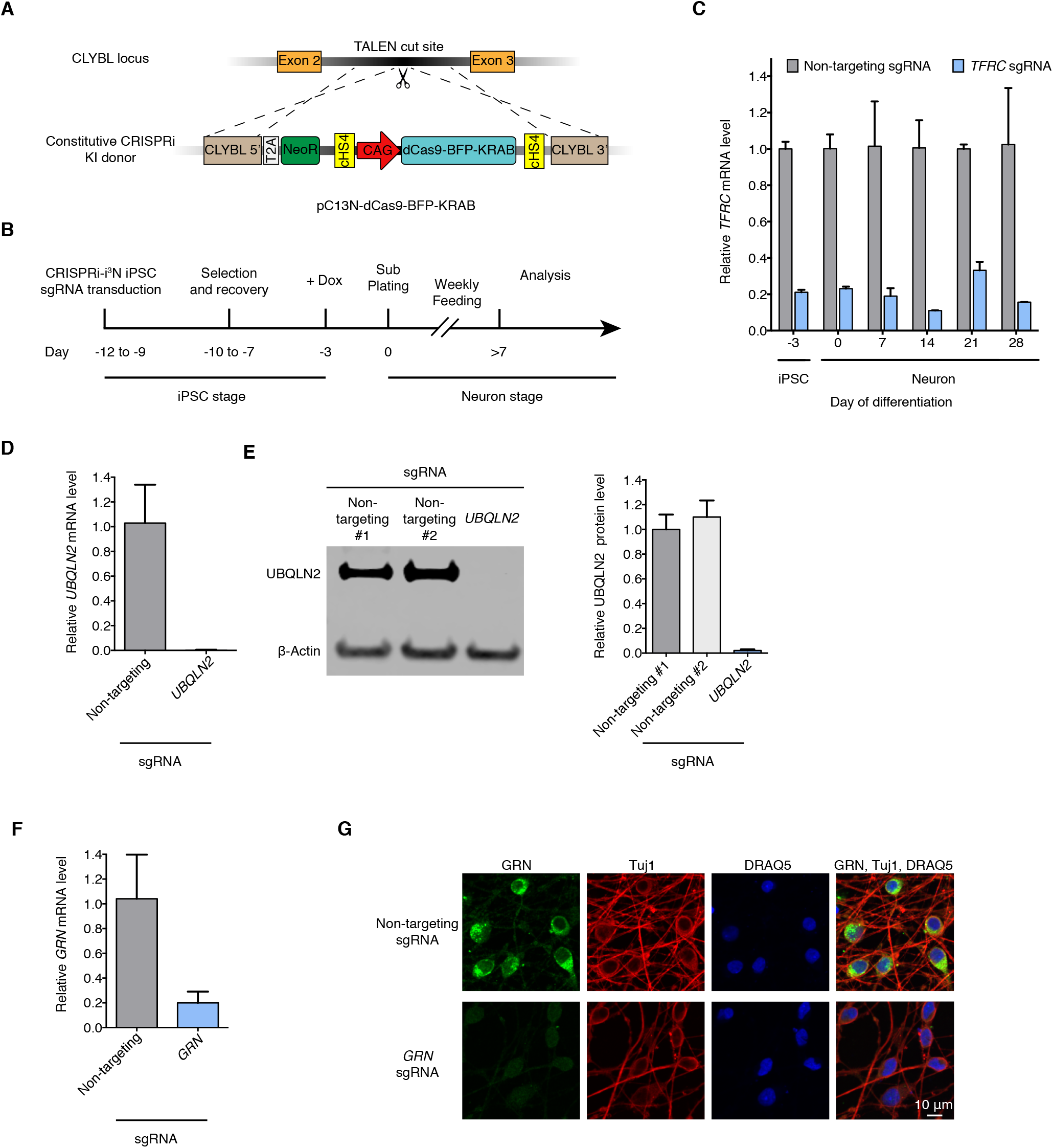
Durable gene knockdown by CRISPR interference in human iPSC-derived neurons. **(A)** Construct pC13N-dCas9-BFP-KRAB for the expression of CRISPRi machinery from the CLYBL safe-harbor locus: catalytically dead Cas9 (dCas9) fused to blue fluorescent protein (BFP) and the KRAB domain, under the control of the constitutive CAG promoter. **(B)** Timeline for sgRNA transduction, selection and recovery, doxycycline-induced neuronal differentiation and functional analysis of CRISPRi-i^3^N iPSCs. **(C)** Knockdown of the transferrin receptor (*TFRC*) in CRISPRi-i^3^N iPSCs and neurons. CRISPRi-i^3^N iPSCs were lentivirally infected with an sgRNA targeting *TFRC* or a non-targeting negative control sgRNA. Neuronal differentiation was induced by addition of doxycycline on Day −3 of the differentiation protocol and plating cells in neuronal medium on Day 0. Cells were harvested at different days for qPCR. After normalizing by *GAPDH* mRNA levels, ratios of *TFRC* mRNA were calculated for cells expressing the TFRC-targeting sgRNA versus the non-targeting sgRNA; mean ± SD (two biological replicates). **(D, E)** Knockdown of ubiquilin 2 (*UBQLN2*) in CRISPRi-i^3^N neurons. CRISPRi-i^3^N neurons infected with *UBQLN2* sgRNA or non-targeting control sgRNA were harvested on Day 11 for qPCR (D) or Western blot (E) to quantify *UBQLN2* knockdown at the mRNA level or protein level, respectively. (D) Relative *UBQLN2* mRNA level was determined by normalizing *UBQLN2* mRNA level by *GAPDH*. Relative *UBQLN2* mRNA was calculated for cells expressing the UBQLN2-targeting sgRNA versus the non-targeting sgRNA; mean ± SD (three biological replicates). (E) *Left*, representative Western blot (Loading control β-Actin). *Right*, quantification of UBQLN2 protein levels normalized by β-Actin for cells with non-targeting sgRNAs or *UBQLN2* sgRNA; mean ± SD (two independent Western blots). **(F,G)** Knockdown of progranulin (*GRN*) in CRISPRi-i^3^N neurons. CRISPRi-i^3^N neurons infected with *GRN* sgRNA or non-targeting control sgRNA were harvested on Day 11 for qPCR (F) or monitored by immunofluorescence (IF) microscopy on Day 5. (G) Relative *GRN* mRNA level normalized by *GAPDH* mRNA. Ratio of relative *GRN* mRNA for cells expressing the GRN-targeting sgRNA versus the non-targeting sgRNA; mean ± SD (three biological replicates). (G) *Top row*, non-targeting negative control sgRNA. *Bottom row*, sgRNA targeting progranulin. Progranulin signal (IF, green), neuronal marker Tuj1 (IF, red) and nuclear counterstain DRAQ5 (blue) are shown.

To validate CRISPRi activity, we transduced these iPSCs with a lentiviral construct expressing an sgRNA targeting the transferrin receptor gene (*TFRC*). Knockdown of *TFRC* mRNA was robust in iPSCs and in i^3^Neurons for several weeks after differentiation (Fig. 1B,C). We also validated knockdown of three additional genes, *UBQLN2* (Fig. 1D,E), *GRN* (Fig. 1F,G) and *CDH2* (Fig. S1B) by qRT-PCR, Western blot, and/or immunofluorescence. Our platform thus enables potent CRISPRi knockdown of endogenous genes in iPSC-derived neurons.

Since CRISPRn-associated DNA damage has been found to be highly toxic to iPSCs (Ihry et al., 2018), we evaluated whether the CRISPRi machinery caused DNA damage in iPSCs or otherwise interfered with neuronal differentiation or activity. We found that expression of CRISPRi machinery and/or sgRNAs did not cause detectable DNA damage (Fig. S1C,D), as expected based on the abrogation of nuclease activity in dCas9, and did not affect neuronal differentiation (Fig. S1E) or activity as evaluated by calcium imaging (Fig. S1F and Movies S1, S2).

We established the CRISPRi-i^3^N system used throughout this study in the background of the well-characterized WTC11 iPSC line (Miyaoka et al., 2014). In addition, we also generated an equivalent line in the NCRM5 iPSC line (Luo et al., 2014) and validated its CRISPRi activity (Fig. S1G).

### A pooled CRISPRi screen reveals neuron-essential genes

We then used this platform to identify cell type-specific genetic modifiers of survival in pooled genetic screen in iPSCs and iPSC-derived neurons (Fig. 2A). We first transduced CRISPRi-i Ni^3^PSCs with our lentiviral sgRNA library H1 (Horlbeck et al., 2016). The H1 library targets 2,325 genes encoding kinases and other proteins representing the “druggable genome” with at least five independent sgRNAs per gene, plus 500 non-targeting control sgRNAs, for a total of 13,025 sgRNAs. Transduced iPSCs were either passaged for 10 days, or differentiated into neurons by doxycycline-induced *Ngn2* expression. Neurons were collected 14, 21 and 28 days post-induction. Frequencies of cells expressing each sgRNA at each time point were determined by next-generation sequencing of the sgRNA-encoding locus. We observed highly correlated sgRNA frequencies between independently cultured experimental replicates (Fig. S2A), supporting the robustness of these measurements.

**Fig. 2.**
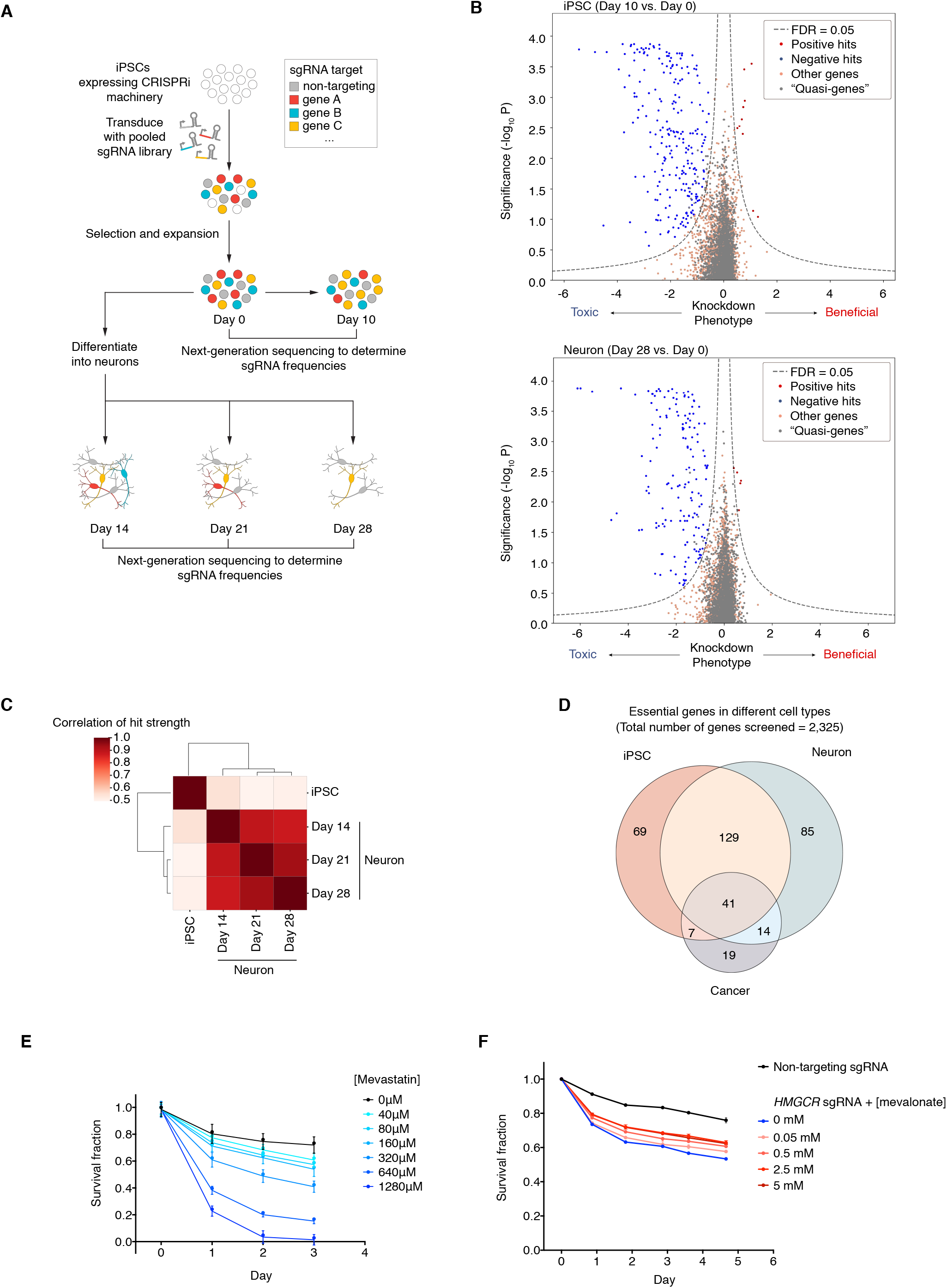
Massively parallel screen for essential genes in iPSCs and iPSC-derived neurons. **(A)** Strategy: CRISPRi-i^3^N iPSCs were transduced with a lentiviral sgRNA library targeting 2,325 genes (kinase and the druggable genome) and passaged as iPSCs or differentiated into glutamatergic neurons. Samples of cell populations were taken at different time points, and frequencies of cells expressing a given sgRNA were determined by next-generation sequencing. **(B)** Volcano plots summarizing knockdown phenotypes and statistical significance (Mann-Whitney U test) for genes targeted in the pooled screen. *Top*, proliferation/survival of iPSCs between Day 0 and Day 10. *Bottom*, survival of iPSC-derived neurons between Day 0 and Day 28. Dashed lines: cutoff for hit genes (FDR = 0.05, see Methods). **(C)** Correlation of hit gene strength (the product of phenotype and –log_10_(P value)) obtained for Day 10 iPSCs, and neurons harvested on Day 14, 21, or 28 post-induction. **(D)** Overlap between essential genes we identified here in iPSCs and neurons, and gold-standard essential genes in cancer cell lines (Hart et al., 2017). **(E)** Survival of neurons without treatment (black) or with various concentrations of Mevastatin (blue) quantified by microscopy; mean ± SD (six replicates). **(F)** Survival of neurons infected with non-targeting sgRNA (black) or *HMGCR* sgRNA (blue) or *HMGCR* sgRNA with various concentrations of mevalonate (pink to red) quantified by microscopy; mean ± SD (six replicates).

To analyze the screen results, we developed a new bioinformatics pipeline, MAGeCK-iNC (MAGeCK including Negative Controls, available at kampmannlab.ucsf.edu/mageck-inc). This pipeline integrates a published method, MAGeCK (Li et al., 2014) with aspects of our previous bioinformatics pipeline (Kampmann et al., 2013, 2014) to take full advantage of the non-targeting control sgRNAs in our library when computing P values (see Methods for details). Based on the depletion or enrichment of sgRNAs targeting specific genes at different time points compared to day 0, we identified hit genes for which knockdown was toxic or beneficial to either iPSCs or neurons at different time points (Fig. 2B, Fig. S2B). We then calculated a knockdown phenotype score and significance P value for each gene (Table S1). The large number of non-targeting sgRNAs in our library enabled us to generate “quasi-genes” from random groupings of non-targeting sgRNAs to empirically estimate a false-discovery rate (FDR) for a given cutoff of hit strength (defined as the product of phenotype score and –log_10_(P value)), see Methods for details. We defined genes passing an FDR < 0.05 as hit genes. For the majority of hit genes, two or three sgRNAs in the library resulted in strong phenotypes (Fig. S2C,D), justifying the use of five sgRNAs/gene in the primary screen library.

Knockdown phenotypes of hit genes were strongly correlated between neurons at different time points, but distinctly less correlated between neurons and iPSCs (Fig. 2C). Next, we compared genes that were essential in iPSCs and/or neurons in our screens with “gold-standard” essential genes that were previously identified through genetic screens in cancer cell lines (Hart et al., 2017). This analysis revealed a shared core set of essential genes, as expected, and additional iPSC-specific and neuron-specific essential genes (Fig. 2D).

Using Gene Set Enrichment Analysis (GSEA) (Mootha et al., 2003; Subramanian et al., 2005), we found enrichment of distinct groups of survival-related genes in neurons compared to iPSCs, such as genes associated with sterol metabolism (Fig. S3A). We validated the strong neuronal dependence on the cholesterol biogenesis pathway pharmacologically using the HMG-CoA reductase inhibitor mevastatin (Fig. 2E) and found that CRISPRi knockdown of HMG-CoA reductase (*HMGCR*) can be partially rescued by supplementing its product mevalonate (Fig. 2F).

We determined expression levels of genes at different time points during neuronal differentiation by Quant-Seq (Data deposited in GEO, GSE124703; the results can be visualized at kampmannlab.ucsf.edu/ineuron-rna-seq). As a group, neuron-essential genes were expressed at significantly higher levels than non-essential genes in iPSC-derived neurons (one-sided Mann-Whitney U test, Fig. S3B). The vast majority of neuron-essential genes were detectable at the transcript level, further supporting the specificity of our screen results.

Intriguingly, we identified several genes that specifically enhanced neuronal survival when knocked down, including *MAP3K12* (encoding dual leucine zipper kinase DLK), *MAPK8* (encoding Jun kinase JNK1), *CDKN1C* (encoding the cyclin-dependent kinase inhibitor p57) and *EIF2AK3* (encoding the eIF2alpha kinase PERK) (Table S1). A pathway involving DLK, JNK and PERK has previously been implicated in neuronal death (Ghosh et al., 2011; Huntwork-Rodriguez et al., 2013; Larhammar et al., 2017; Miller et al., 2009; Pozniak et al., 2013; Watkins et al., 2013; Welsbie et al., 2013), validating our approach.

In summary, our large-scale CRISPRi screen in human iPSC-derived neurons uncovered genes that control the survival of neurons, but not cancer cells or iPSCs, demonstrating the potential of our platform to characterize the biology of differentiated cell types.

### Pooled validation of hit genes

To validate and further characterize hit genes from the primary large-scale screen, we performed a series of secondary screens. For this purpose, we generated a new lentiviral sgRNA plasmid (pMK1334) that enables screens with single-cell RNA-Seq (scRNA-Seq) readouts (based on the CROP-Seq format (Datlinger et al., 2017)), and high-content imaging readouts (expressing a bright, nuclear-targeted BFP) (Fig. 3A). We individually cloned 192 sgRNAs into this plasmid (184 sgRNAs targeting 92 different hit genes with two sgRNAs per gene and eight non-targeting control sgRNAs). Then, to confirm essential genes identified in our primary screen, we pooled these plasmids and conducted a survival-based validation screen (Fig. 3A). Because the library size was small compared to the primary screen, we obtained a high representation of each sgRNA in the validation screen. As in the primary screen, CRISPRi-i^3^N iPSCs transduced with the plasmid pool were either passaged as iPSCs or differentiated into glutamatergic neurons, and then harvested at different time points for next-generation sequencing and calculation of survival phenotypes for each sgRNA (Table S2). We observed a high correlation of raw sgRNA counts between two independently differentiated biological replicates (R^2^ > 0.9, Fig. 3B), supporting the robustness of phenotypes measured in the pooled validation screen. We then compared the results from the validation screen with those from the primary screen. In both iPSCs and neurons, all positive hits and most of the negative hits from the primary screen were confirmed in the validation screen (Fig. 3C). These findings indicate that hits identified in the primary screen are highly reproducible.

**Figure 3.**
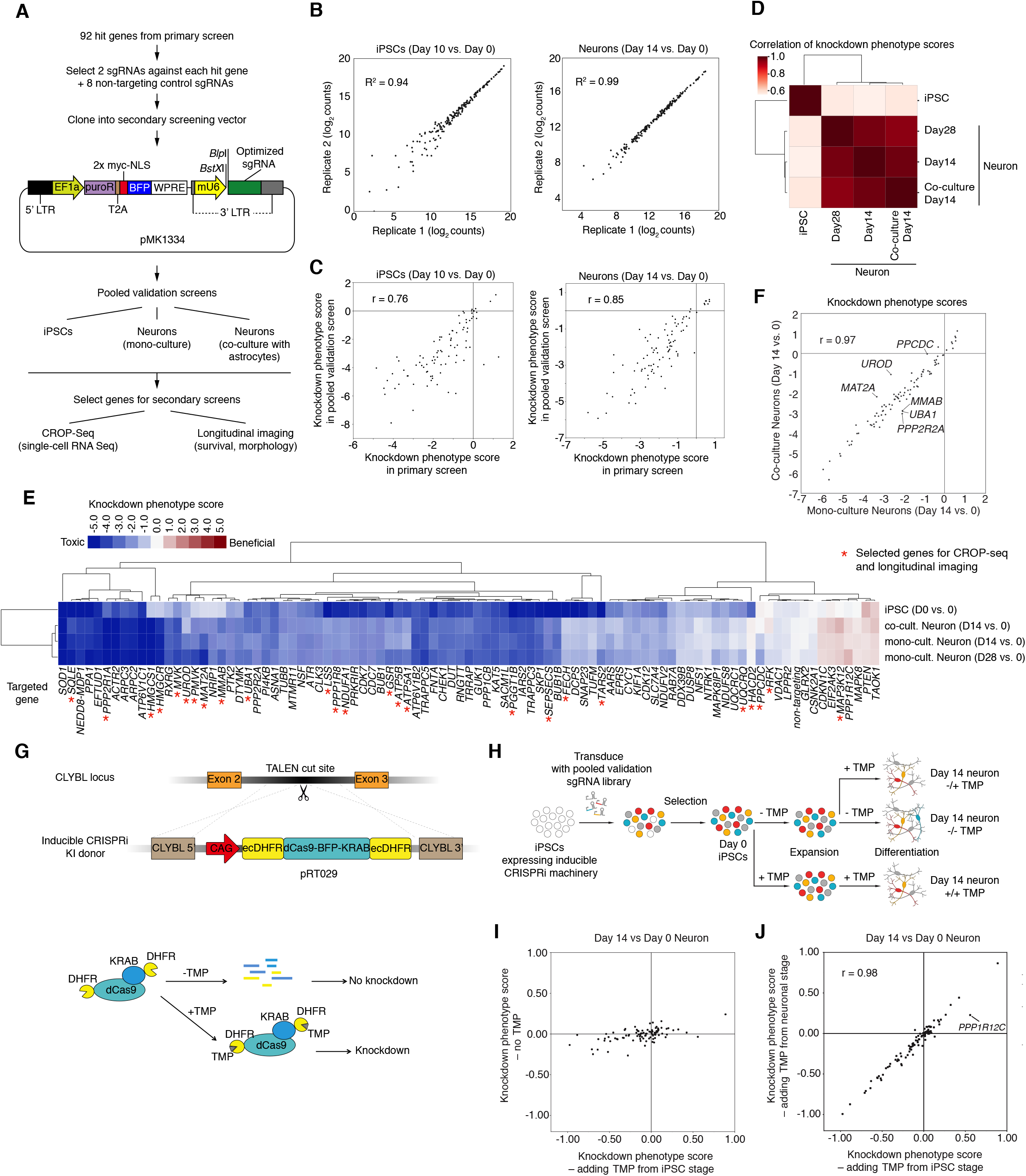
Pooled validation of hit genes from the primary screen. **(A)** Strategy for validation of hit genes. **(B)** Raw counts of sgRNAs from next-generation sequencing for biological replicates of Day 10 iPSCs (left) and Day 14 neurons (right) and coefficients of determination (R^2^), Each dot represents one sgRNA. **(C)** Knockdown phenotype scores from primary screens and validation screens for Day 10 iPSCs (left) and Day 14 neurons (right) and Pearson correlation coefficients (r). Each dot represents one gene. **(D)** Hierarchical clustering of different cell populations from the pooled validation screens based on the pairwise correlations of the knockdown phenotype scores of all genes. **(E)** Heatmap showing knockdown phenotype scores of the genes targeted in the validation screen (columns) in different cell populations (rows). Both genes and cell populations were hierarchically clustered based on Pearson correlation. Red asterisks mark genes selected for secondary screens (CROP-Seq and longitudinal imaging). **(F)** Gene knockdown phenotype scores of Day 14 neurons in monoculture (x-axis) and co-culture with primary mouse astrocytes (y-axis) and Pearson correlation coefficient (r). Each dot represents one gene. Outlier genes, (differences > ± 2 SD from the mean differences) are labeled. **(G)** Strategy for degron-based inducible CRISPRi. Addition of trimethoprim (TMP) stabilizes the DHFR degron-tagged CRISPRi machinery. **(H)** Strategy to test whether hit genes control neuronal survival or earlier processes. **(I)** Knockdown phenotype scores for Day 14 neurons from screens in the inducible CRISPRi iPSCs, comparing populations with TMP added from the iPSC stage (x-axis) to populations without TMP added (y-axis). Each dot represents one gene. **(J)** Knockdown phenotype scores for Day 14 neurons from screens in the inducible CRISPRi iPSCs, comparing populations with TMP added from the iPSC stage (x-axis) to populations with TMP added from the neuronal stage (y-axis) and Pearson correlation coefficient (r). Each dot represents one gene. The outlier gene, *PPP1R12C*, is labeled.

In the brain, many neuronal functions are supported by glial cells, particularly astrocytes. To rule out the possibility that hits from the primary screen were artifacts of an astrocyte-free culture environment, we included an additional condition in the validation screen, in which neurons were co-cultured with primary mouse astrocytes. Neuronal phenotypes in the presence or absence of astrocytes were highly correlated (Fig. 3D,E and Fig. S4A), indicating that the vast majority of the neuron-essential genes we identified are required even in the presence of astrocytes. However, we identified a small number of genes, including *PPCDC, UROD* and *MAT2A*, for which knockdown was less toxic in the presence of astrocytes (Fig. 3F). This suggests that astrocytes may compensate for the loss of function for these genes in neurons. We also identified a small number of other genes, including *MMAB, UBA1* and *PPP2R2A*, for which knockdown was more toxic in the presence of astrocytes (Fig. 3F). These genes may function in pathways affected by crosstalk between neurons and astrocytes.

### Inducible CRISPRi distinguishes neuronal differentiation and survival phenotypes

A caveat of our primary screen is that we introduced the sgRNA library into cells constitutively expressing CRISPRi machinery at the iPSC stage. Therefore, some hit genes detected in the primary screen may play a role in neuronal differentiation rather than neuronal survival. To explore this possibility, we developed a system to allow independent control of neuronal differentiation and CRISPRi activity. We generated inducible CRISPRi constructs by tagging the CRISPRi machinery (dCas9-BFP-KRAB) with dihydrofolate reductase (DHFR) degrons. In the absence of the small molecule trimethoprim (TMP), these DHFR degrons cause proteasomal degradation of fused proteins. Addition of TMP counteracts degradation (Iwamoto et al., 2010). Our initial construct contained a single N-terminal DHFR degron (Fig. S4B), which was insufficient to fully suppress CRISPRi activity in the absence of TMP (Fig. S4C). Therefore, we generated another plasmid (pRT029) with DHFR degrons on both the N- and C-termini of dCas9-BFP-KRAB (Figure 3G). This dual-degron CRISPRi construct was then integrated into the CLYBL locus of i^3^N-iPSCs. In the absence of TMP, the double-degron construct had no CRISPRi activity in iPSCs or neurons (Fig. S4D). TMP addition starting at the iPSC stage resulted in robust CRISPRi activity in iPSCs and neurons (Fig. S4D), and TMP addition starting at the neuronal stage resulted in moderate CRISPRi activity (Fig. S4E). While future optimization of the inducible CRISPRi construct will be necessary, these results indicate that temporal regulation of CRISPRi activity can be achieved in iPSCs and differentiated neurons.

We used the inducible CRISPRi platform to determine if hit genes from our primary screen were related to neuronal survival or differentiation. iPSCs expressing the dual-degron construct were transduced with the pooled validation sgRNA library. Cells were then cultured under three different conditions, including no TMP, TMP added starting at the iPSC stage, and TMP added at the neuronal stage (Fig. 3H). In the population cultured without TMP, none of the sgRNAs showed strong phenotypes compared to cells to which TMP was added at the iPSC stage (Fig. 3I), confirming the tight control of the inducible system. To determine if any of the neuron-essential genes identified in our primary screen were in actuality required for differentiation, we compared neurons in which knockdown was induced either at the iPSC stage or later at the neuronal stage of the protocol. Phenotypes observed in these two conditions were highly correlated (r = 0.98, Fig. 3J), indicating that the vast majority of hits identified from the original screen are indeed essential for neuronal survival, rather than differentiation (Fig. 3H).

Interestingly, there was one exception: sgRNAs targeting *PPP1R12C* were strongly enriched when TMP was added at the iPSC stage, but this phenotype was substantially weaker when TMP was added at the neuron stage. Based on this finding, we hypothesized that these sgRNAs may interfere with neuronal differentiation. Indeed, we observed that two independent sgRNAs targeting *PPP1R12C* each caused continued proliferation instead of neuronal differentiation in a subset of iPSCs (Fig. S4F,G), providing an explanation for the enrichment of cells expressing *PPP1R12C-targeted* sgRNAs in the primary screen. Thus, our inducible CRISPRi approach successfully uncovered a false-positive hit from the primary screen, which affected differentiation as opposed to neuronal survival. Interestingly, the AAVS1 locus, into which the inducible Ngn2 transgene was integrated, resides within the *PPP1R12C* gene. An open question remains as to whether *PPP1R12C* plays a role in neuronal differentiation, or whether sgRNAs directed against *PPP1R12C* interfered with doxycycline-mediated induction of Ngn2.

Taken together, these pooled validation screens confirmed that hits from the primary screen were highly reproducible and that we were able to identify genes specifically essential for neuronal survival.

### CROP-Seq generates mechanistic hypotheses for genes controlling neuronal survival

Recently developed strategies to couple CRISPR screening to scRNA-Seq readouts yield rich, high-dimensional phenotypes from pooled screens (Adamson et al., 2016; Datlinger et al., 2017; Dixit et al., 2016). As a first step towards understanding the mechanisms by which hit genes affect the survival of iPSCs and neurons, we investigated how gene knockdown altered transcriptomes of single cells (Fig. 4A). We selected 27 genes that exemplified different categories of hits based on their pattern of survival phenotypes in iPSCs and neurons (Fig. 3E). A pool of 58 sgRNAs (two sgRNAs targeting each selected gene and four non-targeting control sgRNAs) in the secondary screening plasmid pMK1334 (Fig. 3A) was transduced into CRISPRi-i^3^N iPSCs. We used the 10x Genomics platform to perform scRNA-Seq of ~ 20,000 iPSCs and 20,000 Day 7 neurons. We chose to monitor transcriptomic effects of hit gene knockdown at the early Day 7 time point to capture earlier, gene-specific effects of knockdown, as opposed to later nonspecific effects reflecting toxicity. Transcripts containing sgRNA sequences were further amplified to facilitate sgRNA identity assignment, adapting a previously published strategy (Hill et al., 2018). Following sequencing, transcriptomes and sgRNA identities were mapped to individual cells (Data deposited in GEO, GSE124703). High data quality was evident from the mean reads per cell (~84,000 for iPSCs, ~91,000 for neurons), the median number of genes detected per cell (~5,000 for iPSCs, ~4,600 for neurons) and the number of cells to which a unique sgRNA could be assigned after quality control (~15,000 iPSCs, ~8,400 neurons). Based on the expression of canonical marker genes, we excluded the possibility that gene knockdown interfered with differentiation to glutamatergic neurons (Fig. S5A).

**Figure 4.**
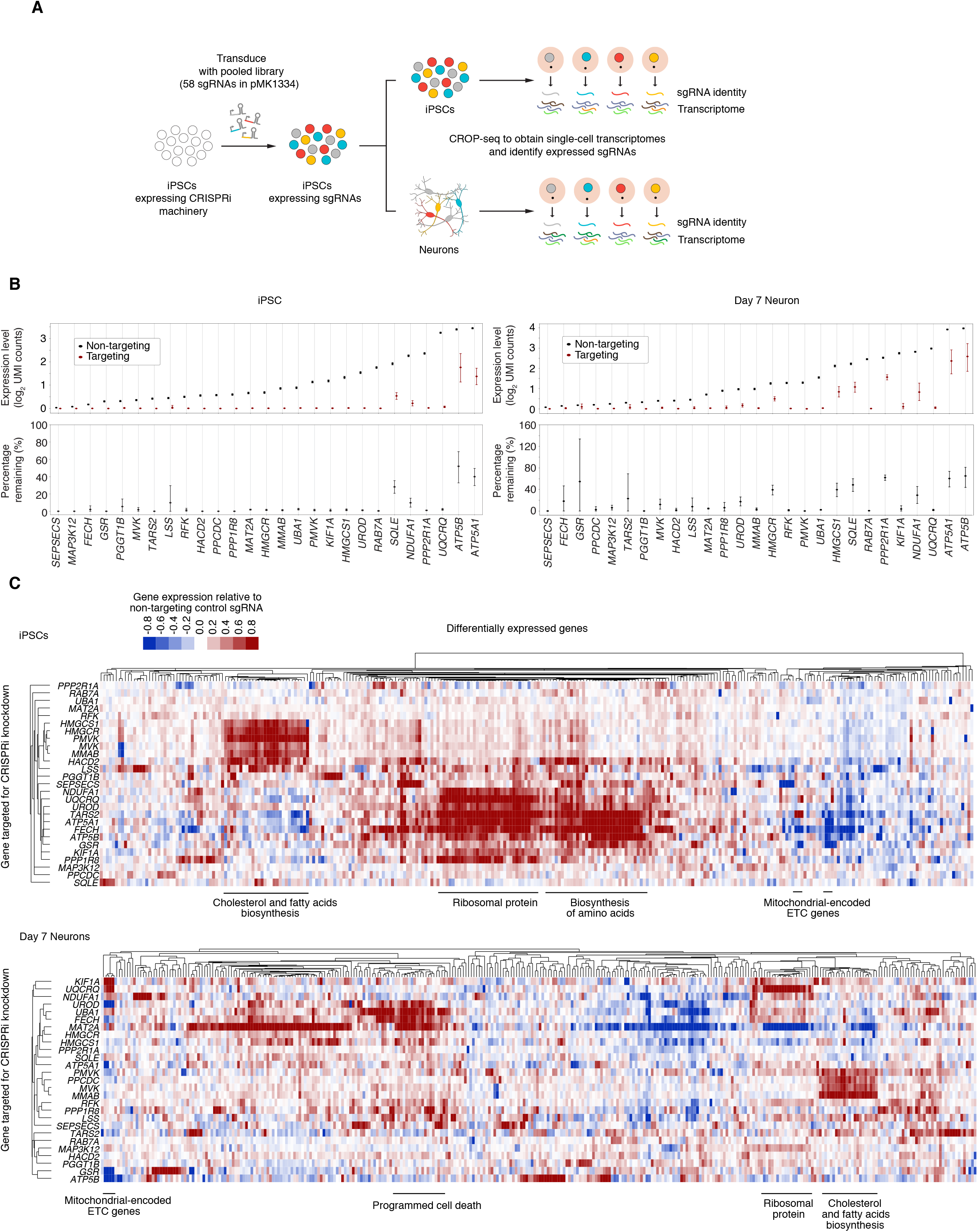
CROP-Seq reveals transcriptome changes in iPSCs and iPSC-derived neurons induced by knockdown of survival-relevant genes. **(A)** Strategy for CROP-Seq experiments. **(B)** On-target knockdown efficiencies in the CROP-Seq screen were quantified for iPSCs (left) and Day 7 neurons (right). For each target gene, the 50% of cells with the strongest on-target knockdown were selected from all cells expressing sgRNAs targeting the gene; average expression of each target gene within these cells is compared to cells with non-targeting control sgRNAs. Error bars: 95% confidence intervals estimated by bootstrapping. **(C)** Changes in gene expression in response to CRISPRi knockdown of genes of interest in iPSCs (top) and Day 7 neurons (bottom). Each row represents one targeted gene; for each targeted gene, the top 20 genes with the most significantly altered expression were selected, and the merged set of these genes is represented by the columns. Rows and columns were clustered hierarchically based on Pearson correlation. Functionally related groups of differentially expressed genes are labeled.

Next, we examined the transcriptomes of groups of cells expressing a given sgRNA (which we refer to as “sgRNA groups”). In both iPSCs and neurons, the two sgRNA groups expressing sgRNAs targeting the same gene tended to form clusters in t-Distributed stochastic neighbor embedding (tSNE) plots (Fig. S5B), confirming that independent sgRNAs targeting the same gene had highly similar phenotypic consequences. The extent of gene knockdown varied across cells within an sgRNA group and between the two sgRNAs targeting the gene. Given that many genes selected for the CROP-Seq screen are essential, it is likely that cells with lower levels of knockdown had a survival advantage and are enriched in the sequenced population. To characterize phenotypes in cells with the most stringent gene knockdown, we took advantage of the single-cell resolution of the CROP-Seq data to select the top 50% of cells with the best on-target knockdown for each gene for further analysis. We refer to this group of cells as the “gene knockdown group”. Compared to cells with non-targeting sgRNAs, the expression levels of the targeted genes in a gene knockdown group were greatly repressed (Fig. 4B). For most genes (24/27 in iPSCs, 18/27 in neurons) knockdown levels of greater than 80% were achieved. Together, these findings further support the robustness of CRISPRi knockdown and of the transcriptomic phenotypes determined by our modified CROP-Seq platform.

To characterize how gene knockdown altered transcriptomes of iPSCs and neurons, we performed differential expression analysis between gene knockdown groups and the negative control group (Table S3). While knockdown of some genes induced the expression of cell-death related genes (including *PDCD2, AEN, GADD45A* and *ATF3*), no generic signature of dying cells dominated the differentially expressed genes. Rather, knockdown of different genes resulted in gene-specific transcriptomic signatures (Fig. 4C). By clustering gene knockdown groups based on the signature of differential gene expression, we found transcriptomic signatures associated with knockdown of functionally related genes (Fig. 4C). For some genes, knockdown resulted in upregulation of functionally related genes. For example, knockdown of genes involved in cholesterol and fatty acid biosynthesis, including *HMGCS1, HMGCR, PMVK, MVK, MMAB*, and *HACD2*, caused induction of other genes in the same pathway (Fig. 4C, Table S3). Thus, pooled CROP-Seq screens can identify and group functionally related genes in human neurons.

The CROP-Seq screen also generated mechanistic hypotheses. For example, knockdown of *MAP3K12* specifically improved neuronal survival. Signaling by the *MAP3K12-encoded* kinase DLK was previously implicated in neuronal death and neurodegeneration (Ghosh et al., 2011; Huntwork-Rodriguez et al., 2013; Larhammar et al., 2017; Miller et al., 2009; Pozniak et al., 2013; Watkins et al., 2013; Welsbie et al., 2013). In our screen, knockdown of *MAP3K12* resulted in coherent changes in neuronal gene expression (Fig. 5A and Table S3). Ribosomal genes and the anti-apoptotic transcription factor Brn3a (encoded by *POU4F1*) were upregulated.

**Figure 5.**
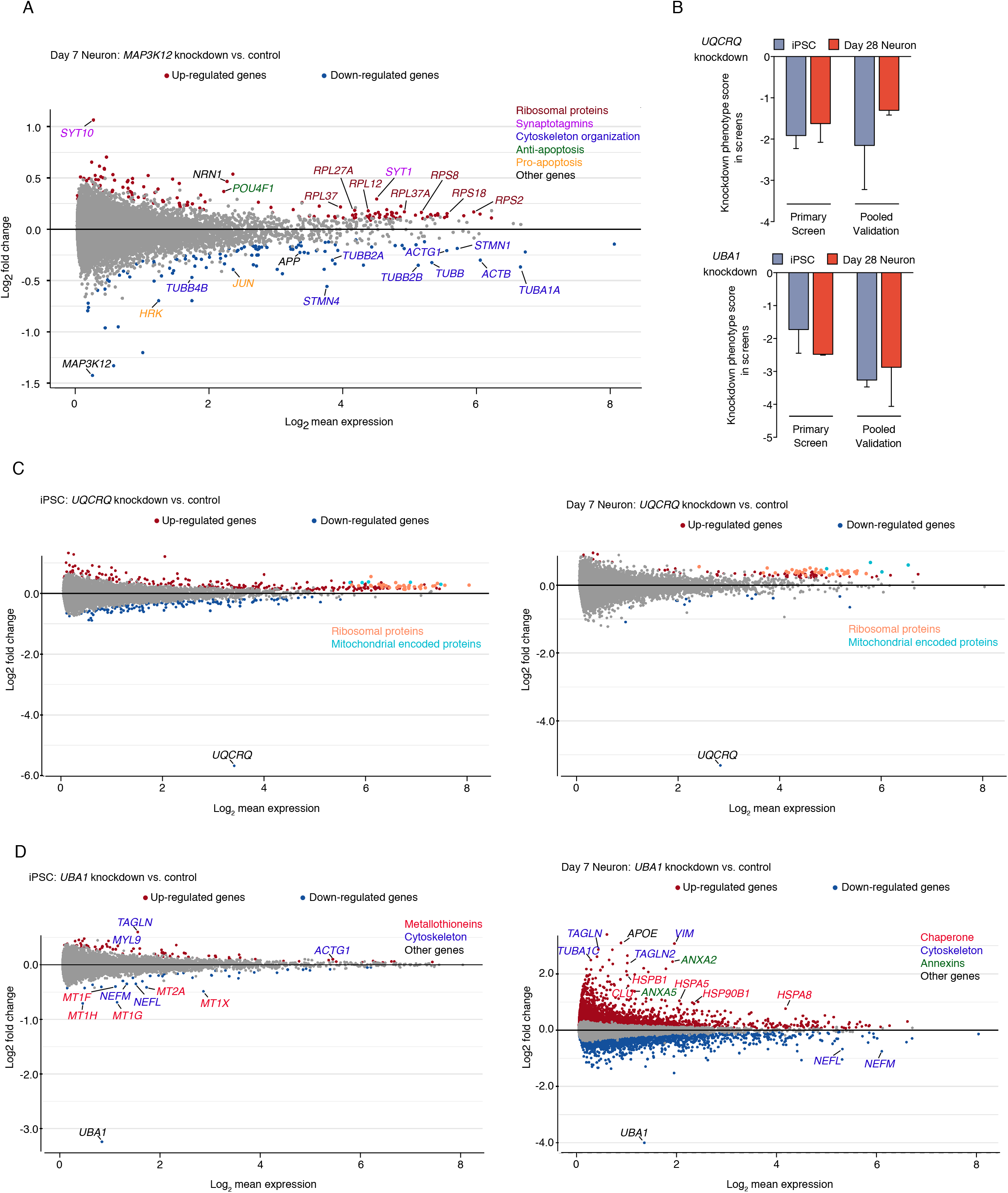
Cell-type specific responses to gene knockdown on the transcriptomic level. **(A)** Changes in transcript levels caused by *MAP3K12* knockdown in neurons from the CROP-Seq screen. Differentially expressed genes (p_adj_ < 0.05) in red (upregulation) or blue (downregulation), or other colors for genes discussed in the main text. **(B)** Knockdown phenotypes of *UQCRQ* (top) and *UBA1* (bottom) in iPSCs and iPSC-derived neurons from the primary and validation screens. Survival phenotypes of 2 sgRNAs targeting the same gene, mean ± SD. **(C,D)** Transcriptomic changes caused by knockdown of *UQCRQ* (C) or *UBA1* (D) in iPSCs and neurons. Differentially expressed genes (p_adj_ < 0.05) in red (upregulation) or blue (downregulation), or other colors for genes discussed in the main text.

Conversely, we observed downregulation of the pro-apoptotic BCL-2 protein Harakiri/DP5 (encoded by *HRK*), the neurodegeneration-associated amyloid precursor protein (*APP*), and the pro-apoptotic transcription factor *JUN*, which is also a downstream signaling target of DLK (Welsbie et al., 2013). Furthermore, *MAP3K12* knockdown caused downregulation of a vast array of proteins involved in cytoskeletal organization, and upregulation of specific synaptotagmins, which act as calcium sensors in synaptic vesicles. These changes in gene expression may relate to the function of DLK in synaptic terminals and its reported role as a neuronal sensor of cytoskeletal damage (Valakh et al., 2015). Lastly, *MAP3K12* knockdown induced expression of neuritin (*NRN1*), a neurotrophic factor associated with synaptic plasticity and neuritogenesis (Cantallops et al., 2000; Javaherian and Cline, 2005; Naeve et al., 1997; Yao et al., 2016). Intriguingly, neuritin levels are decreased in Alzheimer’s Disease patient brains, and overexpression of neuritin was found to be protective in a mouse model of Alzheimer’s Disease (Choi et al., 2014). Thus, CROP-Seq provides a wealth of testable hypotheses for neuroprotective mechanisms and specific effectors downstream of DLK/*MAP3K12* inhibition.

### CROP-Seq reveals neuron-specific transcriptomic consequences of gene knockdown

The results from our parallel CROP-Seq screens in iPSCs and neurons enabled us to compare transcriptomic consequences of gene knockdown across both cell types (Fig. S5C). Interestingly, only a few genes, including *SQLE, MMAB, MVK, UQCRQ*, and *ATP5B*, showed high similarity (similarity score > 0.15) in the transcriptomic changes they induced in iPSCs versus neurons. Knockdown of most genes induced distinct transcriptomic responses in the two cell types. This suggests that either gene knockdown caused different stress states in the two cell types or that gene regulatory networks are wired differently in iPSCs and iPSC-derived neurons.

To further dissect these cell type-specific phenotypes, we ranked genes by the similarity of their knockdown phenotypes in iPSCs and neurons with respect to survival and transcriptomic response (Fig. S5D). For some genes, both survival and transcriptomic phenotypes were similar in iPSCs and neurons. An example for this category of genes is *UQCRQ*, which encodes a component of the mitochondrial complex III in the electron transport chain. *UQCRQ* is essential in both cell types (Fig. 5B), and knockdown of *UQCRQ* had similar transcriptomic consequences in both iPSCs and neurons – upregulation of mitochondrially encoded electron transport chain components and of ribosomal proteins (Fig. 5C, Table S3). Similarly, knockdown of cholesterol and fatty acid biosynthesis genes induced expression of other cholesterol and fatty acid biosynthesis genes in both iPSCs and neurons (Fig. 4C, Table S3).

Interestingly, we also found examples of genes that were essential in both neurons and iPSCs, yet caused substantially different transcriptomic phenotypes when knocked down (Fig. S5D). For example, knockdown of the essential E1 ubiquitin activating enzyme, *UBA1* (Fig. 5B) caused neuron-specific induction of a large number of genes (Fig. 5D, Table S3), including those encoding heat shock proteins (cytosolic chaperones *HSPA8* and *HSPB1* and endoplasmic reticulum chaperones *HSPA5* and *HSP90B1*). This suggests that compromised *UBA1* function triggered a broad proteotoxic stress response in neurons, but not iPSCs, consistent with the role of *UBA1* in several neurodegenerative diseases (Groen and Gillingwater, 2015). Thus, even ubiquitously expressed housekeeping genes can play distinct roles in different cell types.

Lastly, we discovered that some genes differed with respect to both survival and transcriptomic phenotypes in neurons and iPSCs (Fig. S5D). This was expected for genes predominantly expressed in neurons, such as *MAP3K12* (Fig. 5A). However, we also found examples of genes in which knockdown had strikingly different transcriptomic consequences in neurons and iPSCs despite high expression in both cell types. Such a gene is *MAT2A*, encoding methionine adenosyl transferase 2a, which catalyzes the production of the methyl donor S-adenosylmethionine (SAM) from methionine and ATP (Fig. 6A). *MAT2A* is essential in neurons, but not iPSCs (Fig. 6B). Knockdown of *MAT2A* in iPSCs did not substantially affect the expression of any gene other than *MAT2A* itself (Fig. 6C). In neurons, however, knockdown of *MAT2A* caused differential expression of thousands of genes (Fig. 6D, Table S3). Genes downregulated in neurons in response to *MAT2A* knockdown were enriched for neuron-specific functions (Fig. 6E), providing a possible explanation for the neuron-selective toxicity of *MAT2A* knockdown.

**Figure 6.**
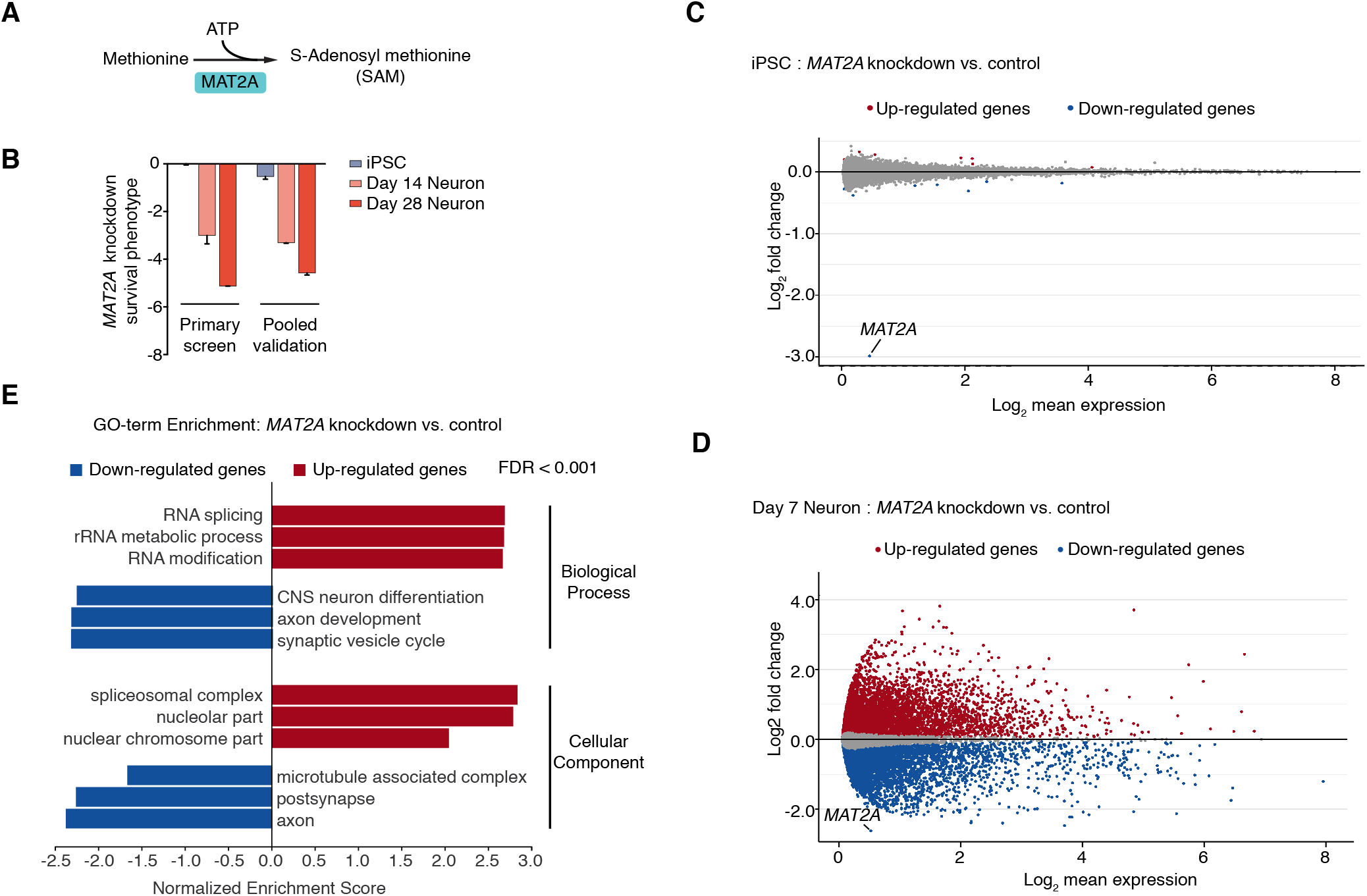
CROP-Seq reveals neuron-specific transcriptomic consequences of *MAT2A* knockdown. **(A)** Methionine adenosyl transferase 2a (*MAT2A*) catalyzes the production of the methyl donor S-adenosylmethionine (SAM) from methionine and ATP. **(B)** *MAT2A* is essential in neurons but not iPSCs. Knockdown phenotypes of *MAT2A* in iPSCs and neurons from the primary and validation screens. Survival phenotypes of 2 sgRNAs targeting *MAT2A*, mean ± SD. **(C,D)** Changes in transcript levels caused by *MAT2A* knockdown in iPSCs (C) and neurons (D) from the CROP-Seq screen. Differentially expressed genes (p_adj_ < 0.05) in red (upregulation) or blue (downregulation). **(E)** Gene Set Enrichment Analysis (GSEA) results for differentially expressed genes in iPSC-derived neurons with *MAT2A* knockdown compared to negative control sgRNAs. Significantly enriched GO terms for Biological Process and Cellular Component are shown.

In summary, results from CROP-Seq screens in iPSCs and iPSC-derived neurons further highlight differences in gene function across the two cell types, provide rich insights into consequences of gene knockdown, and generate mechanistic hypotheses. They further support the idea that it is critically important to study gene function in relevant cell types, even for widely expressed genes.

### An arrayed CRISPRi platform for rich phenotyping by longitudinal imaging

While pooled genetic screens are extremely powerful due to their scalability, many cellular phenotypes cannot be evaluated using a pooled approach. Such phenotypes include morphology, temporal dynamics, electrophysiological properties, and non-cell-autonomous phenotypes. To expand the utility of our screening platform, we therefore optimized an arrayed CRISPRi platform for iPSC-derived neurons.

As a proof-of-concept arrayed screen, we established a longitudinal imaging platform to track the effect of knocking down selected hit genes from our primary screen on neuronal survival and morphology over time. First, we stably expressed cytosolic mScarlet (for neurite tracing) and nuclear-localized mNeon-Green (for survival analysis) in CRISPRi-i^3^N iPSCs. Then, we infected these iPSCs in multi-well plates with lentiviral preparations encoding 48 individual sgRNAs (23 genes selected from the gene set from the CROP-Seq screen targeted by two sgRNAs each, and two non-targeting sgRNAs), followed by puromycin selection and longitudinal imaging of iPSCs, or neuronal differentiation. After three days, we re-plated predifferentiated neurons on 96-well plates alongside similarly prepared cells that did not express sgRNAs or the cytosolic mScarlet marker at a 1:20 ratio to allow more accurate tracing of mScarlet-expressing neurons. These plates were then longitudinally imaged every few days using an automated microscope with a large area of each well imaged at each time point, allowing us to re-image the same populations of neurons over time (Fig. 7A, Movies S3, S4).

**Figure 7.**
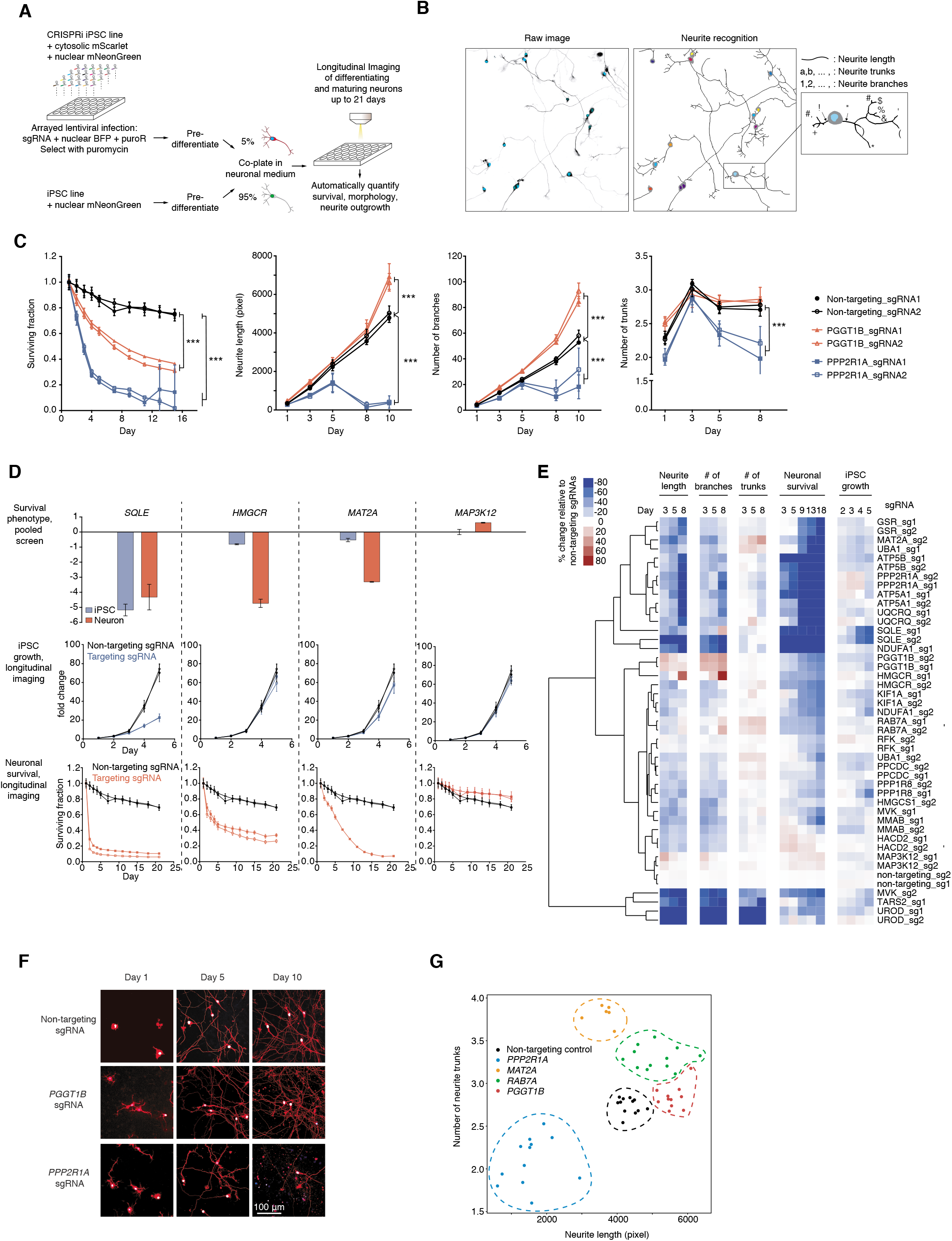
Longitudinal imaging to track the effect of selected hit gene knockdown on iPSC growth, neuronal survival and neurite morphology. **(A)** Strategy for longitudinal imaging for neuronal survival and neurite morphology. **(B)** An example illustrating the image analysis pipeline. A raw image (left) containing sgRNA positive neurons expressing nuclear BFP (cyan) and cytosolic mScarlet (greyscale) were segmented and neurites were recognized (right). Different parameters, including neurite length, number of neurite trunks and number of neurite branches were quantified for individual neurons. Total number of sgRNA positive neurons was quantified for each image to monitor neuronal survival. **(C)** Quantification of knockdown effects of *PGGT1B* and *PPP2R1A* on neuronal survival, neurite length, number of branches and number of trunks. For each sgRNA, mean ± SD of replicate images is shown for each time point. *** significant differences compared to non-targeting sgRNA (P < 0.001, Student’s t-test). **(D)** Examples of hit genes whose survival phenotypes in the pooled screens were validated by longitudinal imaging. *Top*, knockdown phenotypes of *SQLE, HMGCR, MAT2A* and *MAP3K12* in iPSCs and neurons from the validation screens. Average phenotypes of two sgRNAs targeting each gene; error bars represent SD. Growth curves of iPSCs (*middle*) and survival curves of neurons (*bottom*) with non-targeting sgRNAs and sgRNAs targeting *SQLE, HMGCR, MAT2A* or *MAP3K12*. Fold change (for iPSCs, *middle*) or surviving fraction (for neurons, *bottom*) of number of sgRNA-positive cells relative to Day 1 was calculated for each imaging well, mean ± SD for all replicate wells for one sgRNA are shown. **(E)** Changes of iPSC proliferation, neuronal survival and neurite morphology features relative to non-targeting sgRNAs at different time points (columns) induced by knockdown of different genes (rows). Rows were hierarchically clustered based on Pearson correlation. **(F)** Representative images of neurons with *PGGT1B* and *PPP2R1A* knockdown on Days 1, 5 and 10. Nuclear BFP is shown in blue, cytosolic mScarlet is shown in red. Scale bar, 100 μm. **(G)** Effect of gene knockdown on neurite length (x-axis) and number of neurite trunks (y-axis). Each dot indicates the mean measurements of all neurons in one image. Different target genes are shown in different colors, and replicate images for one target gene are grouped by dashed lines in the same colors.

We developed an automated image analysis pipeline to segment neuronal cell bodies and neurites (Fig. 7B). By tracking cell numbers over time, we could measure neuronal survival and iPSC proliferation (Fig. 7C,D). Quantification of survival based on longitudinal imaging was robust across independent experiments (Fig. S6A). Three individual sgRNAs were so toxic that they prevented longitudinal imaging, and were removed from further analysis. As anticipated, the vast majority of sgRNAs that altered survival in pooled screens also altered survival in our arrayed longitudinal survival analysis (Fig. 7D). However, longitudinal imaging provided additional information on the timeline of toxicity caused by knockdown of different genes and revealed gene-specific temporal patterns (Fig. 7D,E).

We then analyzed the effect of gene knockdown on neurite morphology. Our neurite segmentation algorithm extracted multiple morphology metrics, including neurite length, number of neurite trunks and neurite branching (Fig. 7B,C). Our longitudinal imaging approach also enabled us to evaluate adverse effects of the expression of CRISPRi machinery and/or non-targeting sgRNA using highly sensitive readouts. We found that neither CRISPRi machinery nor non-targeting sgRNAs affected neuronal survival (Fig. S6B) or neurite growth (Fig. S6C).

Surprisingly, knockdown of genes that we selected based on their impact on neuronal survival also had distinct effects on neuronal morphology (Fig. 7C,F). Knockdown of the geranylgeranyltransferase *PGGT1B* promoted both neurite growth and branching, consistent with previous findings that protein prenylation inhibits axon growth (Li et al., 2016). Neurite length and the number of neurite trunks were under independent genetic control (Fig. 7G). Taken together, the profile of features extracted from our imaging platform was so information-rich and gene-specific that hierarchical clustering of individual sgRNAs based on these features led to co-clustering of the two sgRNAs targeting a given gene for the majority of genes (Fig. 7E). Conceptually, knockdown phenotypes of specific genes occupy distinct regions in a high-dimensional neuronal morphology space (Fig. 7E,G).

In combination with survival-based and CROP-Seq screens, our arrayed high-content CRISPRi platform will enable the deep characterization of gene function in a plethora of human cell types.

## DISCUSSION

Here, we describe a platform for large-scale, multimodal CRISPRi-based genetic screens in human iPSC-derived neurons. While CRISPR screens in cancer cells and stem cells have revealed numerous biological insights, we reasoned that screens in differentiated, non-cancerous cell types could elucidate novel, cell-type specific gene functions. Indeed, our survival screens uncovered genes that were essential for neurons, but not iPSCs or cancer cells. We also found that knockdown of some broadly-expressed housekeeping genes, such as *UBA1*, caused strikingly distinct transcriptomic phenotypes in neurons compared to iPSCs, consistent with the idea that gene functions can vary across distinct cell types. Lastly, our arrayed screening platform uncovered gene-specific effects on longitudinal survival and neuronal morphology. These proof-of-concept screens have generated a wealth of phenotypic data, which will provide a rich resource for further analysis and the generation of mechanistic hypotheses.

The combination of CRISPRi functional genomics and iPSC-derived neuron technology leverages the strengths of both approaches. Neurons are a highly specialized and disease-relevant cell type, and thus it is crucial to study certain human gene functions in these cells. However, primary human neurons cannot readily be obtained in the quantities and homogeneity needed for large-scale screens. By contrast, human iPSCs have several fundamental qualities ideally suited for screens. They can be made from readily available cells, such as skin fibroblasts or peripheral mononuclear blood cells; they can be genetically engineered and subsequently expanded to generate large numbers of isogenic cells; and they can then be differentiated into a variety of cell types, including specific neuronal subtypes. Differentiation protocols based on induced expression of transcription factors are particularly useful for screens, as they are rapid and yield large numbers of homogeneous neurons. In addition to the Ngn2-driven generation of glutamatergic neurons (Fernandopulle et al., 2018; Wang et al., 2017; Zhang et al., 2013) used here, induced expression of different transcription factors yield other types of neurons, such as motor neurons (Hester et al., 2011; Shi et al., 2018) and inhibitory neurons (Yang et al., 2017). Systematic screens are beginning to uncover additional combinations of transcription factors driving specific neuronal fates (Liu et al., 2018; Tsunemoto et al., 2018). Thus, iPSC technology could be used to generate different neuron types from an isogenic parental cell line, which would facilitate parallel CRISPR screens to dissect neuronal subtype specific gene function. Such screens will address fundamental questions in neuroscience, such as why specific neuronal subtypes are selectively vulnerable in neurodegenerative diseases (Kampmann, 2017). Furthermore, genetic modifier screens in neurons derived from patient iPSCs and isogenic controls have the potential to uncover new disease mechanisms. These discoveries may, in turn, yield new therapeutic strategies to correct cellular defects linked to disease genes. Despite their usefulness, iPSC-derived neurons have limitations – in particular, they do not fully recapitulate all features of mature (or aging) neurons in the human brain. We anticipate that functional genomics approaches, such as ours, may hold the key to improving protocols that lead to ever more faithful models of mature human neurons.

CRISPRi is particularly well suited as a method to study gene function in iPSC-derived neurons, for several reasons. First, it does not cause DNA damage (Fig. S1C,D), and thus lacks the non-specific p53-mediated toxicity observed with CRISPRn approaches in iPSCs and untransformed cells (Haapaniemi et al., 2018; Ihry et al., 2018). Second, it is inducible and reversible (Gilbert et al., 2014), enabling the time-resolved dissection of human gene function. Third, it perturbs gene function via partial knockdown, as opposed to knockout, thereby enabling functional characterization of essential genes, as demonstrated in this study.

There are several areas for further development of our platform. Further optimization of inducible CRISPRi will result in more potent gene repression in mature neurons, leading to increased sensitivity. The standard use of inducible CRISPRi would be preferable in order to initiate gene perturbation in the differentiated cell state, thereby avoiding false-positive phenotypes due to interference with the differentiation process. Also, establishment of our CRISPR activation (CRISPRa) approach in iPSC-derived neurons will enable gain-of-function genetic screens, which yield complementary insights to CRISPRi loss-of-function screens (Gilbert et al., 2014). Finally, using synthetic sgRNAs instead of lentivirus in arrayed CRISPRi screens would substantially increase scalability.

We anticipate that the technology described here can be broadly applied to include additional neuron-relevant readouts, such as multi-electrode arrays (to measure electrophysiological properties) and brain organoids (to assay interactions of neurons with other cell types). However, our technology is not limited to neurons, and should provide a paradigm for investigating the specific biology of numerous other types of differentiated cells. Parallel genetic screens across the full gamut of human cell types cells may systematically uncover context-specific roles of human genes, leading to a deeper mechanistic understanding of how they control human biology and disease.

## ACKNOWLEDGEMENTS

We thank Li Gan, Bruce Conklin, Forbes D. Porter, and Christopher Wassif for support and advice, Kun Leng for sharing unpublished constructs, Andrew Hill for advice on CROP-Seq, Kun Leng, Emmy Li and Rene Sit for advice on scRNA-Seq, Sami Barmada for advice on longitudinal neuron survival analysis, Marcin Szczot for advice on the calcium imaging experiments, Nina Dräger for gift of primary astrocytes, Ling Hao and Ryan Prestil for protein and RNA samples, Stewart Humble for advice on qRT-PCR and iPSC genome engineering, Eric Chow and Derek Bogdanoff for next-generation sequencing, the CTRC (Combined Technical Research Core/NIDCR) at the NIH for FACS, Avi Samelson for comments on this manuscript, and members of the Kampmann and Ward Labs for discussions.

This research was supported by the Intramural Research Program of the NIH/NINDS, an NIH Director’s New Innovator Award (NIH/NIGMS DP2 GM119139 to M.K.), NIH/NIA grants (R01 AG062359 and R56 AG057528 to M.K.), the NINDS Tau Center Without Walls (NIH/NINDS U54 NS100717 to M.K.), an Allen Distinguished Investigator Award to M.K. (Paul G. Allen Family Foundation, to M.K.), a Chan-Zuckerberg Biohub Investigator Award (to M.K.), and a Tau Consortium Investigator Award (Rainwater Charitable Foundation, to M.K.).

## AUTHOR CONTRIBUTIONS

R.T., M.A.G., C.H.L., M.E.W. and M.K. contributed to the study’s overall conception, design and interpretation, and wrote the manuscript and created the Figures with input from the other authors. R.T. designed and conducted the validation screens and CROP-Seq screens and designed and conducted computational analyses for all screens (including survival screens, CROP-Seq, and longitudinal imaging screens), with guidance from M.K. M.A.G. designed and conducted imaging screens with guidance from M.E.W. and M.K. C.H.L. established CRISPRi in iPSC-derived neurons and designed and conducted the primary screens with guidance from R.T. and M.K. C.H.L. also conducted and analyzed Quant-Seq experiments and created the iNeuronRNASeq web application. M.A.G., M.T.L., M.S.F., and R.P. designed, generated and characterized constructs and cell lines, with guidance from M.E.W. and M.K. J.Y.H. and D.N. generated and characterized constructs and cell lines, with guidance from R.T. and M.K. A.V.P. contributed to validation experiments and M.A. optimized FACS for iPSC line construction.

## DECLARATIONS OF INTEREST

M.K. has filed a patent application related to CRISPRi and CRISPRa screening (PCT/US15/ 40449) and serves on the Scientific Advisory Board of Engine Biosciences.

## STAR METHODS

### CONTACT FOR REAGENT AND RESOURCE SHARING

*Further information and requests for resources and reagents should be directed to and will be fulfilled by the Lead Contact, Martin Kampmann (*).

### EXPERIMENTAL MODEL AND SUBJECT DETAILS

#### Human iPSCs

Human iPSCs (male WTC11 background (Miyaoka et al., 2014) unless otherwise noted; male NCRM5 (Luo et al., 2014) background in Fig. S1G) were cultured in Essential 8 Medium (Gibco/Thermo Fisher Scientific; Cat. No. A1517001) on BioLite Cell Culture Treated Dishes (Thermo Fisher Scientific; assorted Cat. No.) coated with Growth Factor Reduced, Phenol Red-Free, LDEV-Free Matrigel Basement Membrane Matrix (Corning; Cat. No. 356231) diluted 1:100 in Knockout DMEM (Gibco/Thermo Fisher Scientific; Cat. No. 10829-018). Briefly, Essential 8 Medium was replaced every other day or every day once 50% confluent. When 80-90% confluent, cells were passaged, which entailed the following: aspirating media, washing with DPBS, incubating with StemPro Accutase Cell Dissociation Reagent (Gibco/Thermo Fisher Scientific; Cat. No. A11105-01) at 37°C for 7 minutes, diluting Accutase 1:5 in DPBS, collecting in conicals, centrifuging at 200g for 5 minutes, aspirating supernatant, resuspending in Essential 8 Medium supplemented with 10nM Y-27632 dihydrochloride ROCK inhibitor (Tocris; Cat. No. 125410), counting, and plating onto Matrigel-coated plates at desired number.

#### Human iPSC-derived neurons

Human iPSCs engineered to express mNGN2 under a doxycycline-inducible system in the AAVS1 safe harbor locus were used for the differentiation protocol below. iPSCs were released and centrifuged as above, and pelleted cells were resuspended in N2 Pre-Differentiation Medium containing the following: Knockout DMEM/F12 (Gibco/Thermo Fisher Scientific; Cat. No. 12660-012) as the base, 1X MEM Non-Essential Amino Acids (Gibco/Thermo Fisher Scientific; Cat. No. 11140-050), 1X N2 Supplement (Gibco/Thermo Fisher Scientific; Cat. No. 17502-048), 10ng/mL NT-3 (PeproTech; Cat. No. 450-03), 10ng/mL BDNF (PeproTech; Cat. No. 450-02), 1μg/mL Mouse Laminin (Thermo Fisher Scientific; Cat. No. 23017-015), 10nM ROCK inhibitor, and 2μg/mL doxycycline hydrochloride (Sigma-Aldrich; Cat. No. D3447-500MG) to induce expression of mNGN2. iPSCs were counted and plated at 7 x 10^5^ cells per Matrigel-coated well of a 6-well plate in 2mL of N2 Pre-Differentiation Medium, or at 4 x 10^6^ cells per Matrigel-coated 10-cm dish in 12mL of medium, for three days. After three days, hereafter Day 0, pre-differentiated cells were released and centrifuged as above, and pelleted cells were resuspended in Classic Neuronal Medium containing the following: half DMEM/F12 (Gibco/Thermo Fisher Scientific; Cat. No. 11320-033) and half Neurobasal-A (Gibco/Thermo Fisher Scientific; Cat. No. 10888-022) as the base, 1X MEM Non-Essential Amino Acids, 0.5X GlutaMAX Supplement (Gibco/Thermo Fisher Scientific; Cat. No. 35050-061), 0.5X N2 Supplment, 0.5X B27 Supplement (Gibco/Thermo Fisher Scientific; Cat. No. 17504-044), 10ng/mL NT-3, 10ng/mL BDNF, 1μg/mL Mouse Laminin, and 2μg/mL doxycycline hydrochloride. Pre-differentiated cells were subsequently counted and plated plated at 2 x 10^5^ cells per well of a BioCoat Poly-D-Lysine 12-well plate (Corning; Cat. No. 356470) in 1mL of Classic Neuronal Medium, or at 7.5 x 10^6^ cells per BioCoat Poly-D-Lysine 10-cm dish (Corning; Cat. No. 356469) in 10mL medium. On Day 7, half of the medium was removed and an equal volume of fresh Classic Neuronal Medium without doxycycline was added. On Day 14, half of the medium was removed and twice that volume of fresh medium without doxycycline was added. On Day 21, one-third of the medium was removed and twice that volume of fresh medium without doxycycline was added. On Day 28 and each week after, one-third of the medium was removed and an equal volume of fresh medium without doxycycline was added.

For the longitudinal imaging screens, updated media formulations were used for neuronal differentiation and culture. During the three days of pre-differentiation, we used Induction Medium containing the following: Knockout DMEM/F12 (Gibco/Thermo Fisher Scientific; Cat. No. 12660-012) as the base, 1X GlutaMAX Supplement (Gibco/Thermo Fisher Scientific; Cat. No. 35050-061), 1X MEM Non-Essential Amino Acids (Gibco/Thermo Fisher Scientific; Cat. No. 11140-050), 1X N2 Supplement (Gibco/Thermo Fisher Scientific; Cat. No. 17502-048), 10nM ROCK inhibitor, and 2ug/mL doxycycline (Sigma #D9891). Differentiated neurons were cultured in Cortical Neuron Culture Medium containing the following: BrainPhys Neuronal Medium (STEMCELL Technologies #05790) or BrainPhys without Phenol Red (STEMCELL Technologies #05791) as the base, 1X B27 Supplement (Gibco/Thermo Fisher Scientific; Cat. No. 17504-044), 10ng/mL NT-3 (PeproTech; Cat. No. 450-03), 10ng/mL BDNF (PeproTech; Cat. No. 450-02), 1ug/mL Mouse Laminin (R&D Systems #3446-005-01), and optionally 2μg/mL doxycycline.

#### Primary mouse astrocytes

Primary mouse astrocytes were isolated from two P1 mouse pups and cultured in T75 in DMEM + 10% FBS. One day after plating neurons, astrocytes were dissociated by trypsin, washed by PBS to remove any remaining FBS and centrifuged at 200g for 5 minutes. The pelleted astrocytes were resuspended in the Classic Neuronal Medium and plated onto the neuronal culture at a 1:5 astrocytes to neurons ratio. Media changes were performed as indicated above for neuronal culture. Once astrocytes were confluent, 2 μM final concentration of AraC was added to the culture.

### METHOD DETAILS

#### Molecular Cloning

The CLYBL-targeting constitutive CRISPRi vector pC13N-dCas9-BFP-KRAB was obtained by sub-cloning dCas9-BFP-KRAB from plasmid pHR-SFFV-dCas9-BFP-KRAB downstream of a CAG promoter in the CLYBL-targeting pC13N-iCAG.copGFP vector via BsrGI and AgeI digestion, thus replacing copGFP and generating the plasmid. pHR-SFFV-dCas9-BFP-KRAB was a gift from Stanley Qi & Jonathan Weissman, Addgene plasmid # 46911; http://n2t.net/addgene:46911; RRID:Addgene_46911 (Gilbert et al., 2013) and pC13N-iCAG.copGFP was a gift from Jizhong Zou (Addgene plasmid # 66578; http://n2t.net/addgene:66578; RRID:Addgene_66578 (Cerbini et al., 2015)).

The AAVS1-targeting constitutive CRISPRi vector pMTL3 was obtained by inserting a gene block (gBlock, IDT Technologies) encoding BFP-KRAB into pAAVS1-NC-CRISPRi to create a C-terminal fusion with dCas9, and replacing the neomycin resistance marker with a puromycin resistance marker. pAAVS1-NC-CRISPRi (Gen3) was a gift from Bruce Conklin (Addgene plasmid # 73499; http://n2t.net/addgene:73499; RRID:Addgene_73499 (Mandegar et al., 2016)). The degron controlled version was generated by inserting a gene-block encoding *E. coli* dihydrofolate reductase (ecDHFR)–derived degrons with the R12Y, G67S, and Y100I mutations (Iwamoto et al., 2010) to generate an N-terminal fusion with the CRISPRi machinery.

The CLYBL-targeted inducible CRISPRi construct pRT029 was generated by sub-cloning gene blocks encoding *E. coli* dihydrofolate reductase (ecDHFR)–derived degrons with the R12Y, G67S, and Y100I mutations in the first degron and R12H, N18T and A19V in the second degron (Iwamoto et al., 2010) to generate both an N-terminal and a C-terminal in-frame fusion with dCas9-BFP-KRAB in pC13N-dCas9-BFP-KRAB.

The secondary screening vector pMK1334 was generated as follows: The PpuMI – SnaBI fragment of CROPseq-Guide-Puro was replaced with a gene block encoding the mU6-BstXI-BlpI-optimized sgRNA backbone fragment from our sgRNA vector pCRISPRia-v2 (Addgene plasmid # 84832; http://n2t.net/addgene:84832; RRID:Addgene_84832 (Horlbeck et al., 2016)) to obtain pMK1332. CROPseq-Guide-Puro was a gift from Christoph Bock (Addgene plasmid # 86708; http://n2t.net/addgene:86708; RRID:Addgene_86708 (Datlinger et al., 2017)). Next, the RsrII + PflMI fragment from pMK1332 was replaced by the RsrII + PflMI fragment from pCRISPRia-v2 to introduce tagBFP, creating pMK1333. Last, tagBFP was replaced by a gene block encoding 2xmycNLS-tagBFP2 to obtain pMK1334.

The mNeon-Green-NLS vector (H53) was generated by sub-cloning an EF1α promoter and mNeon-Green with two SV40-NLS into the pMK1333 vector via XhoI and EcoRI digestion, thus replacing the mU6 promoter, the original EF1α promoter, and the original fluorophore. The mScarlet vector (I2) was generated by sub-cloning mScarlet downstream of an EF1α promoter in the H53 vector via BmtI and EcoRI digestion, thus replacing mNeon-Green-NLS.

The GCaMP6m vector (I1) was generated by sub-cloning GCaMP6m downstream of an EF1α promoter in the H53 vector via BmtI and EcoRI digestion, thus replacing mNeon-Green-NLS.

Vector maps are available at kampmannlab.ucsf.edu/resources, and plasmids will be shared on Addgene.

#### CRISPRi iPS cell line generation

WTC11 iPSCs harboring a single-copy of doxycycline-inducible mouse NGN2 at the AAVS1 locus (Wang et al., 2017)(Fernandopulle et al., 2018) were used as the parental iPSC line for further genetic engineering. iPSCs were transfected with pC13N-dCas9-BFP-KRAB and TALENS targeting the human CLYBL intragenic safe harbor locus (between exons 2 and 3) (pZT-C13-R1 and pZT-C13-L1, Addgene #62196, #62197) using DNA In-Stem (VitaScientific). After 14 days, BFP-positive iPSCs were isolated via FACS sorting, and individualized cells were plated in a serial dilution series to enable isolation of individual clones under direct visualization with an inverted microscope (EtaLuma LS 620) in a tissue culture hood via manual scraping. Clones with heterozygous integration of dCas9-BFP-KRAB (determined using PCR genotyping) were used for further testing. Karyotype testing (Cell Line Genetics) was normal for the clonal line used for further experiments in this study, which we termed CRISPRi-i^3^N iPSCs. Similarly, we generated the inducible CRISPRi iPSC line by using pRT029 as a donor plasmid, instead of pC13N-dCas9-BFP-KRAB.

NCRM5 iPSCs (Luo et al., 2014) were used as a second parental iPSC line for genetic engineering. iPSCs were transfected using Lipofectamine Stem (ThermoFisher #STEM00003) with Alt-R S.p. HiFi Cas9 Nuclease V3 (Integrated DNA Technologies #1081060), a custom sgRNA targeting the human CLYBL intragenic safe harbor locus (between exons 2 and 3) from Synthego (sequence ATGTTGGAAGGATGAGGAAA), and the following plasmids: pC13N-dCas9-BFP-KRAB, CLYBL-TO-hNGN2-BSD-mApple (Addgene #124229) and pCE-mp53DD (Okita et al., 2013)(Addgene #41856). The following day, iPSCs were individualized and plated in a serial dilution series to enable isolation of individual clones under direct visualization with an inverted microscope (EtaLuma LS 620) in a tissue culture hood via manual scraping. While iPSCs were plated in the serial dilution series, 20 μM blasticidin (Gemini Bio-Products #400-165P) was added to the culture medium for 1-4 days to select for clones with successful integration of TO-hNGN2-BSD-mApple. Clones with heterozygous integration of dCas9-BFP-KRAB and TO-hNGN2-BSD-mApple (determined using PCR genotyping) were used for further testing, including functional CRISPRi activity (verified by GRN immunocytochemistry) and neuronal differentiation (verified visually and with TUJ1 immunocytochemistry).

#### Dissociation of neurons

Papain (Worthington; Code: PAP2; Cat. No.LK003178) was resuspended in 1X Hanks’ Balanced Salt Solution (Corning; Cat. No. 21-022-CV) to 20U/mL and warmed at 37°C for 10 minutes. Magnesium chloride was added at 5mM and DNase (Worthington; Code: DPRF; Cat. No. LS006333) was added at 5ug/mL immediately before use. Culture medium was aspirated and human iPSC-derived neurons were washed with DPBS. The papain, magnesium chloride, and DNase solution was added at 250μL per well of a 12-well plate or at 2mL per 10-cm dish and incubated at 37°C for 10 minutes. This dissociation solution was quenched in 5 volumes of DMEM (Gibco/Thermo Fisher Scientific; Cat. No. 10313-039) supplemented with 10% fetal bovine serum for each volume of dissociation solution, and the resulting solution was used to detach and transfer the sheet of cells to the appropriate collection tube format. For DNA, RNA, or protein extraction, the neuron sheet was centrifuged at 200g for 3 minutes, the supernatant was carefully removed with a P1000 pipette, and the pellet was snap frozen in liquid nitrogen. For flow cytometry analysis, the neuron sheet was triturated 10-15 times and centrifuged at 200g for 10 minutes, the supernatant was carefully removed with a P1000 pipette, and the pellet was resuspended in staining solution.

#### Quantification of knockdown by qPCR

To quantify *TFRC* or *CDH2* knockdown, human iPSCs or neuron cell pellets were thawed on ice, and total RNA was extracted using the Quick-RNA Miniprep Kit (Zymo; Cat. No. R1054). An input of 100ng RNA was used to synthesize cDNA with the SuperScript III First-Strand Synthesis System (Invitrogen; Cat. No. 18080-051). Samples were prepared for qPCR in technical duplicates in 15μL reaction volumes using SensiFAST SYBR Lo-ROX 2X master mix (Bioline; Cat. No. BIO-94005), custom qPCR primers from Integrated DNA Technologies used at a final concentration of 0.4uM, and cDNA (prepared above) diluted at 1:2 or 1:100 for the target or housekeeping gene, respectively. Quantitative real-time PCR was performed on an Agilent Mx3005P QPCR System with the following Fast 2-Step protocol: 1) 95°C for 2 minutes; 2) 95°C for 5 seconds (denaturation); 3) 60°C for 15 seconds (annealing/extension); 4) repeat steps 2 and 3 for a total of 40 cycles; 5) ramp from 55°C to 95°C to establish melting curve. Expression fold changes were calculated using the ΔΔCt method.

To quantify *UBQLN2* or *GRN* knockdown, total RNA was extracted from Day 11 CRISPRi-i3N neurons on 12-well plates using the Direct-zol 96 RNA Kit (Zymo #R2055). Samples were prepared for RT-qPCR in technical and biological triplicates in 10 μL reaction volumes using the iTaq Universal Probes One-Step Kit (Bio-Rad #172-5141). The following PrimePCR Probe Assays from Bio-Rad were diluted 1:20: GAPDH (Unique Assay ID qHsaCEP0041396), UBQLN2 (qHsaCEP0055207), and GRN (qHsaCEP0057821). Quantitative real-time PCR was performed on a QuantStudio 6 Flex Real-time PCR System (ThermoFisher #4485691). Expression fold changes were calculated using the ΔΔCt method.

#### Quantification of knockdown by Western Blot

To quantify protein level knockdown of *UBQLN2* by *UBQLN2* sgRNA in CRISPRi-i^3^N neurons, neurons with 2 different non-targeting sgRNAs or *UBQLN2* sgRNA were lysed and 20ug of total protein from each lysate was loaded into a NuPAGE 4-12% Bis-Tris Gel (Invitrogen, Cat# NP0336BOX). Subsequently, the gel was transferred onto a nitrocellulose membrane, which was then blocked by Odyssey^®^ Blocking Buffer (PBS) (LI-COR, Cat#927-50000), followed by overnight incubation with primary antibodies at 4 degree. The primary antibodies used were Mouse monoclonal anti-β-Actin (8H10D10) (Cell Signaling Technology, Cat#3700) and Rabbit monoclonal anti-UBQLN2 (D7R2Z) (Cell Signaling Technology, Cat#85509). After incubation, the membrane was washed three times with TBST and then incubated with secondary antibodies (LI-COR Cat# 926-32211 and 926-68070) at room temperature for 1hr. The membrane was then washed 3 times with TBST and once with TBS and imaged on the Odyssey Fc Imaging system (LI-COR Cat# 2800). Digital images were processed and analyzed using ImageJ.

#### Immunocytochemistry

To evaluate N-cadherin (*CDH2*) knockdown, pre-differentiated neurons were plated in Classic Neuronal Medium with doxycycline on day 0 at 4 x 10^4^ cells per well on sterilized, Matrigel-coated 12mm diameter round glass coverslips (Ted Pella Inc; Cat. No. 26023) placed in 24-well plates. One day later, primary rat cortical astrocytes (gift from Li Gan) were plated in Classic Neuronal Medium with doxycycline on the same coverslips (co-culture) at 8 x 10^3^ cells per well. On days 7 and 14, half of the medium was removed and an equal volume of fresh Classic Neuronal Medium without doxycycline was added. On day 18, culture medium was aspirated from each well and cells were subsequently washed with DPBS. Cells were then fixed with 4% paraformaldehyde, which was prepared by diluting 16% paraformaldehyde (Electron Microscopy Sciences; Cat. No. 15710) 1:4 in DPBS, at room temperature for 15 minutes. Paraformaldehyde was removed with a P1000 pipette and collected for proper disposal, and coverslips were washed three times with DPBS for 5 minutes each. Cells were blocked with 2.5ug/mL Mouse BD Fc Block (BD Biosciences; Cat. No. 553141) at room temperature for 15 minutes and subsequently incubated with 1uL mouse IgG1k anti-human CD325 antibody conjugated to APC (BioLegend; Cat. No. 350808) in 50uL of Fc Block at room temperature for 45 minutes. Coverslips were then washed once with DPBS for 5 minutes, incubated with 1ug/mL Hoechst 33342 (Thermo Fisher Scientific; Cat. No. H3570) diluted in DPBS at room temperature for 10 minutes, and then washed twice more with DPBS for 5 minutes each. One drop of Aqua Poly Mount (Polysciences; Cat. No. 18606)

To evaluate progranulin (*GRN*) knockdown, pre-differentiated neurons were plated in Cortical Neuron Culture Medium with doxycycline on Day 0 at 3.0 x 10^5^ cells per well on poly-L-ornithine-coated 8-well glass-bottom slides (Ibidi #80827). On day 5, culture medium was aspirated from each well and cells were subsequently washed with PBS. Cells were fixed with 4% paraformaldehyde, which was prepared by diluting 16% paraformaldehyde (Electron Microscopy Sciences; Cat. No. 15710) 1:4 in PBS, at room temperature for 30 minutes. Paraformaldehyde was removed with a P1000 pipette and collected for proper disposal, and slides were washed three times with PBS. Cells were blocked with 3% donkey serum with 0.1% saponin in PBS at room temperature for one hour and subsequently incubated with goat anti-human progranulin antibody diluted 1:3000 (R&D Systems #AF2420) and mouse anti-human TUJ1 antibody (BioLegend #801201) diluted 1:1000 in blocking buffer at 4°C overnight. Slides were then washed three times with PBS and incubated with donkey anti-goat IgG conjugated to AF-488 or AF-647 (Jackson ImmunoResearch #705-545-147 or #705-605-147) and donkey anti-mouse IgG conjugated to RRX or AF-488 (Jackson ImmunoResearch #715-295-151 or #715-545-151) diluted 1:2000 in blocking buffer at room temperature for one hour. Slides were again washed three times with PBS, and incubated with 5 uM DRAQ5 (Thermo Fisher Scientific #62251) in blocking buffer at room temperature for 30 minutes.

To evaluate neuronal differentiation, pre-differentiated neurons were resuspended in Cortical Neuron Culture Medium with doxycycline (2 ug/uL), then plated on poly-L-ornithine-coated 96-well culture dishes (Perkin Elmer #6055308) at a density of 7.5×10^4^ cells/well (n=6 wells per sgRNA). On day 14, culture medium was aspirated from each well on one set of plates and cells were subsequently washed with PBS. Cells were fixed with 4% paraformaldehyde, which was prepared by diluting 16% paraformaldehyde (Electron Microscopy Sciences; Cat. No. 15710) 1:4 in PBS, at room temperature for 15 minutes. Paraformaldehyde was removed and collected for proper disposal, and the plate was washed three times with PBS. Cells were blocked with 5% donkey serum and 0.1% Triton X-100 in PBS at room temperature for 2.5 hours and subsequently incubated with guinea pig anti-NeuN antibody (MilliporeSigma #ABN90) diluted 1:1000 in blocking buffer at 4°C overnight. The plate was then washed three times with PBS and incubated with goat anti-guinea pig IgG conjugated to AF-647 (ThermoFisher Scientific #A-21450) diluted 1:1000 in blocking buffer at room temperature for 2.5 hours. Plates were again washed three times with PBS. Cells without sgRNA were then incubated with 4 uM Hoechst 33342 (Thermo #62249) for 30 minutes at room temperature, and subsequently washed three times with PBS. i^3^N iPSCs plated on Ibidi slides were stained alongside the neurons for comparison. For NeuN quantification, stained neurons were imaged with a spinning disk confocal microscope with a motorized stage (Nikon Eclipse Ti), controlled using Nikon Elements software. A 20X objective was used to acquire a series of 36 slightly overlapping images within each well followed by image stitching.

#### DNA damage assay

CRISPRi-i^3^N iPSCs were infected by non-targeting sgRNA or 2 different MAT2A sgRNAs for 48hrs on Matrigel-coated 96-well plates. Cells with no treatment and with 1uM Etoposide treatment for 6hrs were used as negative and positive controls, respectively. These cells were fixed by 4% paraformaldehyde for 15 mins followed by permeabilization by 0.1% Triton for 10 mins. After that, the cells were blocked with 5% goat serum and 0.1% Triton X-100 in PBS at room temperature for 1 hour and subsequently incubated with mouse anti-H2AX pS139 antibody (Millipore #05-636) diluted in blocking buffer at a final concentration of 2 μg/ml overnight at 4 degree. Following incubation, the cells were washed three times with PBS and incubated with goat anti-mouse IgG conjugated to Alexa Fluor 488 (Abcam, Cat#ab150113) at room temperature for 1hr. Cells were then washed three times with PBS. Untreated cells and Etoposide treated cells were incubated with 1uM Hoechst 33342 (Thermo #62249) for 30 minutes at room temperature. For quantification, stained iPSCs were imaged using an InCell 6000 (GE Cat# 28-9938-51) at 60X and H2AX foci were quantified using CellProfiler (Carpenter et al., 2006).

#### Calcium Imaging

i^3^N iPSCs and CRISPRi-i^3^N iPSCs were lentivirally transduced with the GCaMP6m vector (I1). These polyclonal iPSCs were passaged and plated at a density of 5.0×10^4^ cells/well in Matrigel-coated 6-well culture plates. Shortly afterwards, half of the wells with iPSCs were transduced with lentivirus containing non-targeting sgRNA. The following day, media was changed to E8 + RI. Two days after infection, the media was changed to E8 + 12 ug/mL puromycin (Sigma #P9620-10ML) to select for transduced cells.

Following selection for 3-4 additional days, the iPSCs were passaged into fresh Matrigel-coated 6-well culture plates at a high density and allowed to differentiate in Induction Medium with doxycycline (2 ug/mL) for 3 days, with daily media changes.

Following the 3 days of differentiation, these partially-differentiated neurons were passaged and resuspended in Cortical Neuron Culture Medium with doxycycline (2 ug/uL), then plated on poly-L-ornithine-coated 96-well culture dishes (Perkin Elmer #6055308) at a density of 7.5×10^4^ cells/well (n=6 wells per sgRNA). For the remainder of the experiment before imaging, half of the culture medium was removed and an equal volume of fresh medium was added three times per week. On day 28, half of the culture medium was aspirated from each well. Neurons were imaged with a spinning disk confocal microscope with a 37 °C heated chamber and a motorized stage (Nikon Eclipse Ti), controlled using Nikon Elements software. A 20X objective was used to acquire 30-second movies of one field per well (n = 4-6 wells) at approximately 5 frames/second. Following the initial acquisition, culture medium containing CNQX (Tocris #1045) at a final concentration of 50 uM was added to each well. Beginning one minute after CNQX addition, the same fields were imaged as previously.

#### Primary CRISPRi screen

The CRISPRi v2 H1 library with top 5 sgRNAs per gene (Horlbeck et al., 2016) was packaged into lentivirus for transduction of iPSCs as follows. Two 15-cm dishes were each seeded with 8 x 106 HEK293T cells in 20 mL DMEM complete (basal medium supplemented with 10% FBS and 1% penicillin/streptomycin). The next day, H1 library transfection mix was prepared in the following manner: 10ug H1 library plasmid and 10ug third generation packaging mix (1:1:1 mix of the three plasmids) were diluted into 2mL Opti-MEM I Reduced Serum Medium (Gibco; Cat. No. 31985070); 250uL Lipofectamine 2000 Transfection Reagent (Invitrogen; Cat. No. 11668027) was diluted into 2mL Opti-MEM and incubated at room temperature for 5 minutes; the diluted DNA solution was added to the diluted Lipofectamine solution, inverted several times to mix, and incubated at room temperature for 15 minutes. Following incubation, half of the transfection solution was gently added dropwise to each 15-cm dish with HEK293T cells, and the plates were briefly and gently moved in a figure-eight pattern to mix. Eight hours later, the Lipofectamine-containing media on each dish was carefully aspirated and replaced with 20mL DMEM complete supplemented with 40uL ViralBoost (Alstem; Cat. No. VB100; diluted 1:500 in media). Two days later, HEK293T media (approximately 40mL) was transferred to a 50mL conical and centrifuged at room temperature for 10 minutes at 300g to pellet cell debris. The supernatant was carefully transferred to a syringe fitted with a 0.45um filter in order to filter the virus-containing solution into a new 50mL conical. Approximately 10mL of cold Lentivirus Precipitation Solution (Alstem; Cat. No. VC100) was added to this filtered solution, which was then mixed well and stored at 4°C for 48 hours. Following incubation, the solution was centrifuged at 4°C for 30 minutes at 1,500g, and the supernatant was decanted. A second centrifugation at 4°C for 5 minutes at 1,500g was performed, and the remaining supernatant was removed with a P1000 pipette. The virus-containing pellet was resuspended in 20mL Essential 8 iPSC medium with ROCK inhibitor.

For infection with the H1 library, two T175 Matrigel-coated flasks were each seeded with 2 x 10^7^ CRISPRi-i^3^ N iPSCs in 10mL of the virus-containing medium and left in the tissue culture hood for 15 minutes to allow even distribution and attachment before moving to the incubator. Six hours later, an additional 15mL of Essential 8 medium with ROCK inhibitor was added to each flask without removing the virus-containing medium. The next day, we performed a complete media change on all flasks, adding 35mL Essential 8 medium with ROCK inhibitor to allow the cells to recover and proliferate. One day later, we released the cells and seeded four T175 Matrigel-coated flasks each with 1 x 10^7^ cells in 20mL Essential 8 medium with ROCK inhibitor, which was the medium volume and formulation used for puromycin treatment to enrich sgRNA-expressing cells. The initial MOI, quantified as the fraction of BFP-positive cells by flow cytometry, was ~15%, corresponding to a library representation of ~450 cells per library element. Puromycin treatment proceeded in the following manner: two days with 0.8ug/mL puromycin, followed by two days with 1ug/mL puromycin. At the end of treatment, cells were assessed by flow cytometry (83% expressed high levels of BFP) and seeded for the iPSC and neuronal survival screens, which are described below.

For the iPSC growth-based screen, two T175 Matrigel-coated flasks were each seeded with 1 x 10^7^ cells in 20mL Essential 8 medium with ROCK inhibitor (timepoint t_0_), corresponding to a library representation of ~1,200 cells per library element. Approximately 2 x 10^7^ t_0_ cells were also snap frozen in liquid nitrogen for downstream sample preparation to represent the Day 0 sample, corresponding to a library representation of ~1,200 cells per library element. Media was replaced on day two (t_2_), omitting ROCK inhibitor. Cells were released on day three (t_3_), and each replicate was seeded into two new T175 Matrigel-coated flasks with 1 x 10^7^ cells each in 20mL Essential 8 medium with ROCK inhibitor. Media was replaced on day five (t_5_), omitting ROCK inhibitor. Cells were released on day six (t_6_), cells within the same replicate were mixed across flasks, and each replicate was seeded into two new T175 Matrigel-coated flasks with 1 x 10 cells each in 20mL Essential 8 medium with ROCK inhibitor. Media was replaced on days eight (t_8_) and nine (t_9_), omitting ROCK inhibitor. Cells were released on day ten (t_10_), cells within the same replicate were mixed across flasks, and 4 x 10^7^ cells from each replicate were snap frozen for downstream sample preparation, corresponding to a library representation of ~2,500 cells per library element

For the neuronal survival screen, twelve 10-cm Matrigel-coated dishes were each seeded with 4 x 10^6^ iPSCs in N2 Pre-Differentiation Medium (day-3) and differentiated as previously described. However, an additional full media change (10mL) was performed on Day 4 to remove cellular debris that started to appear. On Days 14, 21, and 28, dead (floating cells) were removed and live (adherent) cells from two 10-cm dishes were harvested per replicate per timepoint. Since neuronal death occurred over time, the estimated library representation for these time points was ~410 cells/library element on Day 14, ~380 cells/library element on Day 21, and ~330 cells/library element on Day 28. Adherent cells were released by papain as previously described, and pelleted cells were snap frozen for downstream sample preparation. Genomic DNA was extracted with the NucleoSpin Blood L or XL kits (Macherey Nagel; Cat. No. 740954.20 or 740950.10, respectively) and samples were prepared for sequencing on an Illumina HiSeq-4000 based on previously described protocols (Gilbert et al., 2014; Kampmann et al., 2014).

#### Pooled validation screen

192 sgRNAs, including 184 sgRNAs targeting 92 selected hit genes from the primary screen (two sgRNAs per gene) and 8 non-targeting control sgRNAs, were individually cloned into the secondary screening vector pMK1334 and verified by Sanger sequencing. The plasmid was pooled and lentivirus was produced as for the Primary Screen. CRISPRi-i^3^N iPSCs were transduced with the pool at 70% MOI (quantified as fraction of BFP-positive cells by flow cytometry) and were transduced cells selected by 1 ug/ml of puromycin to obtain a population of cells that was ~85% BFP-positive. Following 3 days of expansion, approximately 2 million of these cells were harvested as Day 0 sample (corresponding to a library representation of ~9,000 cells/library element) and the rest of cells were cultured as iPSCs (as described in ‘Human iPS cell culture’) or differentiated into glutamatergic neurons (as described in ‘Human neuronal culture’). For the iPSC growth screen, iPSCs were cultured in E8 medium with daily medium change in two T25 flasks as duplicates and were passaged every 2-3 days till Day 10. Approximately 2 million of Day 10 iPSCs from each replicate were harvested, corresponding to a library representation of ~9,000 cells/library element. For the mono-culture neuronal screen, 10 million of pre-differentiated neurons were plated in one Poly-D-Lysine coated 15-cm dish (Corning; Cat. No. 354550). Two replicate dishes of neurons were cultured in Classic Neuronal Medium as described in ‘Human neuronal culture’. Live neurons were harvested on Day 14 and Day 28 neurons as described for the primary screens; the library representation was ~35,000 cells / library element on Day 14 and ~28,000 cells / library element on Day 28. For the co-culture neuronal screen, 1.5 million of primary mouse astrocytes were added into one Poly-D-Lysine coated 15-cm dish containing 7.5 million neurons. Two replicate dishes of neurons in co-culture were cultured as described in ‘Astrocyte co-culture’. Day 14 neurons from each replicate of co-culture experiment were harvested, the library representation was ~50,000 neurons/library element. Genomic DNA was isolated from all harvested samples using a commercial kit (Macherey Nagel; NucleoSpin^®^ Blood). The sgRNA-encoding region were then amplified and sequenced as in the Primary Screen.

#### CROP-Seq

CRISPRi-i^3^N iPSCs were infected with a pool of selected sgRNAs in the CROP-Seq vector pMK1334 at a low multiplicity of infection to minimize double infection. After puromycin selection and expansion, cells were either passaged as iPSCs or differentiated into neurons. Approximately 20,000 iPSCs and 20,000 day 7 i^3^Neurons were captured by the 10X Chromium Controller using Chromium Single Cell 3’ Library & Gel Bead Kit v2 (10X Genomics; Cat. No. 120267) with 10,000 input cells per lane. Sample prep was performed according to protocol, holding 10-30 ng full-length cDNA for sgRNA-enrichment PCR.

To facilitate sgRNA assignment, sgRNA-containing transcripts were additionally amplified by hemi-nested PCR reactions by adapting a previously published approach (Hill et al., 2018). Briefly, in the first PCR reaction, 15ng of full-length cDNA was used as template and Enrichemnt_PCR_1_For and Enrichemnt_PCR_1_Rev were used as primers. PCR product was cleaned up by 1.0x SPRI beads (SPRIselect; BECKMAN COULTER; Cat. No. B23317) and 1ng cleaned product was input into the second PCR reaction using Enrichemnt_PCR_2_For and Enrichemnt_PCR_2_Rev as primers. Following 1.0x SPRI beads clean up, 1 ng of the PCR product from the second PCR reaction was used as template in the final PCR, in which reverse primer Enrichemnt_PCR_2_Rev and a forward primer, Enrichemnt_PCR_3_For, containing an i7 index, were used as primers. All PCR reactions were carried out for 18 cycles using KAPA HiFi polymerase (KAPA HiFi HotStart ReadyMix (2X); Cat. No. KK2602) with annealing temperature at 62 degree and 15 seconds extension per cycle. The sgRNA-enrichment libraries were separately indexed and sequenced as spike-ins alongside the whole-transcriptome scRNA-Seq libraries using NovaSeq 6000 using the following configuration:

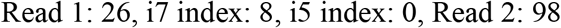

#### Quant-Seq

Neurons cultured in 12-well plates were released with papain, pelleted, and snap frozen on days 0, 14, 21, 28, and 35 in technical duplicates (approximately 2 x 10^5^ cells each) per timepoint. RNA was extracted using the Quick-RNA Miniprep Kit (Zymo; Cat. No. R1054), and RNA concentrations were determined with the Qubit RNA HS Assay Kit (Invitrogen; Cat. No. Q32855) on a Qubit 2.0 Fluorometer (Invitrogen; Cat. No. Q32866). mRNA-Seq libraries were prepared from an input of 184ng total RNA in 5uL using the QuantSeq 3’ mRNA-Seq Library Prep Kit FWD for Illumina (Lexogen; Cat. No. 015). Briefly, oligodT hybridization enabled mRNA-selective reverse transcription. The original RNA template was then degraded, and second strand cDNA synthesis was achieved by random priming and extension by DNA polymerase. Samples were subsequently subjected to magnetic bead-based purification, followed by library amplification with indexed flow-cell adapters (14 PCR cycles) and another round of magnetic bead-based purification. mRNA-Seq library concentrations (mean of 1.01 ± 0.275 ng/uL) were measured with the Qubit dsDNA HS Assay Kit (Invitrogen; Cat. No. Q32851) on a Qubit 2.0 Fluorometer. Library fragment-length distributions (mean of 371 ± 16.1 bp) were quantified with the High Sensitivity DNA Kit (Agilent; Cat. No. 5067-4626) on a 2100 Bioanalyzer Instrument (Agilent; Cat. No. G2939BA). Molar concentrations for each sample were approximated from Qubit concentration (ng/uL) and mean fragment-length (bp) measurements using the following formula:

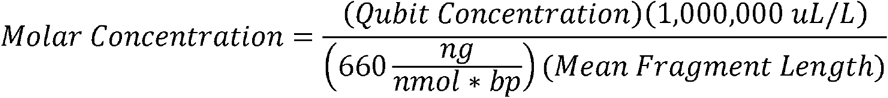

All libraries were diluted to 2.72nM for equimolar representation in the final, pooled sample. Single-end sequencing was performed, generating reads toward the poly(A) tail.

#### Longitudinal CRISPRi-i^3^Neuron imaging screen

CRISPRi-i^3^N iPSCs were transduced with lentivirus expressing mNeonGreen-NLS and FACS-sorted for the brightest green population. These polyclonal cells will be referred to as nuclear-green CRISPRi-i^3^N iPSCs. Subsequently, a fraction of these iPSCs were transduced with lentivirus expressing cytosolic mScarlet and FACS sorted for the brightest red and green cells. These polyclonal iPSCs will be referred to as the nuclear-green/cytosolic red CRISPRi-i^3^N iPSCs.

The arrayed sgRNAs in the pMK1334 vector were packaged into lentivirus for transduction of iPSCs as follows: 6-well plates coated with poly-D-lysine were seeded with 2.5 x 10^6^ HEK293T cells per well in 1.5 mL of DMEM complete (basal medium supplemented with 10% FBS) each. The next day, the arrayed transfection mixes were prepared in the following manner: 1.2ug sgRNA plasmid and 1.2 ug packaging mix (0.8ug psPAX2, 0.3ug pMD2G, 0.1ug pAdVantage), along with 5 uL P3000 reagent (ThermoFisher Scientific # L3000015) were diluted into 150 uL Opti-MEM I Reduced Serum Medium (Gibco; Cat. No. 31985070); 3.75 uL Lipofectamine 3000 Transfection Reagent (ThermoFisher Scientific # L3000015) was diluted into 150 uL Opti-MEM and incubated at room temperature for 5 minutes; the diluted Lipofectamine solution was added to the diluted DNA solution, flicked to mix, and incubated at room temperature for 20 minutes. Following incubation, the transfection solutions were gently added dropwise to each well with HEK293T cells, and the plates were briefly and gently moved in a figure-eight pattern to mix. The following day, the Lipofectamine-containing media on each well was carefully aspirated and replaced with 3mL DMEM complete supplemented with 6uL ViralBoost (Alstem; Cat. No. VB100; diluted 1:500 in media). Three days later, HEK293T media from each well was transferred to one well each of two 2mL deep 96-well dishes (USA Scientific #1896-2800) and centrifuged at 4°C for 30 minutes at 3428g to pellet cell debris. Viral supernatant was stored at 4°C.

For functional titering of the lentivirus, nuclear-green+ CRISPRi-i^3^N iPSCs were passaged and plated at a density of 1.0 x 10^4^ cells/well in Matrigel-coated 96-well culture dishes. Following adherence of iPSCs, 75 uL of each viral supernatant was added to one well, and a series of half-volume dilutions was performed for a total of four dilutions. The following day, the culture medium containing the lentivirus was carefully aspirated and replaced with fresh medium. Three days after infection, iPSCs were imaged with a spinning disk confocal microscope with a motorized stage (Nikon Eclipse Ti), controlled using Nikon Elements software. A 20X objective was used to acquire a series of 36 slightly overlapping images within each well followed by image stitching. The ratio of cells infected with lentivirus was quantified via Nikon Elements software as the number of green nuclei with blue signal above an intensity threshold divided by the total number of green nuclei. The volumes of viral supernatant used in all subsequent infections were adjusted based on the differences between infection ratios.

Nuclear-green+ cytosolic red+ CRISPRi-i^3^N iPSCs were passaged and plated at a density of 2.5×10^4^ cells/well in Matrigel-coated 12-well culture plates. Shortly afterwards, the iPSCs were transduced with lentivirus containing individual sgRNAs. The following day, media was changed to E8 + RI. Two days after infection, the media was changed to E8 + puromycin (12 ug/mL) to select for transduced cells.

Following selection for 3-4 additional days, the iPSCs were passaged into fresh Matrigel-coated 12-well culture plates at a high density and allowed to differentiate in Induction Medium with doxycycline (2 ug/mL) for 3 days, with daily media changes. Concurrently, the uninfected nuclear-green CRISPRi-i^3^N iPSCs were differentiated alongside the infected nuclear-green+blue/cytosolic red CRISPRi-i^3^N iPSCs.

Following the 3 days of differentiation, these partially-differentiated neurons were passaged and resuspended in Cortical Neuron Culture Medium with doxycycline (2 ug/uL), then plated on poly-L-ornithine-coated 96-well culture dishes (Perkin Elmer #6055308) at a density of 5.0×10^4^ cells/well (n=6 wells per sgRNA for most, 3-5 for some). The nuclear-green+blue/cytosolic red CRISPRi-i^3^Neurons were spiked in at a density of 1:20 with the nuclear-green CRISPRi-i^3^Neurons to facilitate tracing of neurites while maintaining trophic support of higher-density neuron cultures. Following plating, we waited for adherence of neurons before imaging for the first time. For the remainder of each longitudinal imaging experiment, half of the culture medium was removed and an equal volume of fresh medium was added three times per week.

For each timepoint, CRISPRi-i^3^Neurons were imaged with a spinning disk confocal microscope with a motorized stage (Nikon Eclipse Ti), controlled using Nikon Elements software. A 20X objective was used to acquire a series of 25 slightly overlapping images within each well followed by image stitching. Between imaging sessions, plates were incubated in a traditional water-jacketed 5% CO_2_ incubator at 37°C.

#### Longitudinal iPSC imaging screen

Nuclear-green CRISPRi-i^3^N iPSCs were passaged and plated in Matrigel-coated 96-well culture dishes at a density of 1,000 cells/well. Following adherence, iPSCs were transduced with lentivirus (same preparation as for the longitudinal neuronal imaging) containing individual sgRNAs (n=3 wells per sgRNA). The following day, media was changed to E8 + ROCK inhibitor. Starting two days after infection, iPSCs were imaged with a spinning disk confocal microscope with a motorized stage (Nikon Eclipse Ti), controlled using Nikon Elements software. A 20X objective was used to acquire a series of 36 slightly overlapping images within each well followed by image stitching. Between imaging sessions, plates were incubated in a traditional water-jacketed 5% CO_2_ incubator at 37 °C.

#### Longitudinal imaging for CRISPRi toxicity

i^3^N iPSCs and inducible CRISPRi-i^3^N iPSCs were transduced with lentivirus expressing only mNeonGreen-NLS, and/or with lentivirus expressing cytosolic mScarlet. These polyclonal iPSC groups will be referred to as the nuclear-green iPSCs and nuclear-green/cytosolic red iPSCs, respectively. Nuclear-green/cytosolic red iPSCs were passaged and plated at a density of 5.0×10^4^ cells/well in Matrigel-coated 6-well culture plates. Shortly afterwards, half of the wells with iPSCs were transduced with lentivirus containing non-targeting sgRNA. The following day, media was changed to E8 + RI. Two days after infection, the media was changed to E8 + puromycin (12 ug/mL) to select for transduced cells. Following selection for 3-4 additional days, the iPSCs were passaged into fresh Matrigel-coated 12-well culture plates at a high density and allowed to differentiate in Induction Medium with doxycycline (2 ug/mL) for 3 days, with daily media changes. Concurrently, the uninfected nuclear-green iPSCs were differentiated alongside the nuclear-green/cytosolic red iPSCs. Following the 3 days of differentiation, these partially-differentiated neurons were passaged and resuspended in Cortical Neuron Culture Medium with doxycycline (2 ug/uL), then plated on poly-L-ornithine-coated 96-well culture dishes (Perkin Elmer #6055308) at a density of 5.0×10^4^ cells/well (n=6 wells per sgRNA). The inducible CRISPRi-i^3^N neurons were plated in two groups, with or without 20 uM TMP (Sigma #92131-5G) in the culture medium. The nuclear-green/cytosolic red neurons were spiked in at a density of 1:20 with the nuclear-green neurons to facilitate tracing of neurites while maintaining trophic support of higher-density neuron cultures. Following plating, we waited for adherence of neurons before imaging for the first time. For the remainder of each longitudinal imaging experiment, half of the culture medium was removed and an equal volume of fresh medium was added three times per week. For each timepoint, neurons were imaged with a spinning disk confocal microscope with a motorized stage (Nikon Eclipse Ti), controlled using Nikon Elements software. A 20X objective was used to acquire a series of 25 slightly overlapping images within each well followed by image stitching. Between imaging sessions, plates were incubated in a traditional water-jacketed 5% CO2 incubator at 37 °C.

#### Pharmacological validation of *HMGCR* phenotype

CRISPRi-i^3^N cells infected with non-targeting control sgRNA or *HMGCR* sgRNA were seeded into 96- or 384-well plates on Day 0 into Brainphys media containing mevastatin (compactin, Sigma #M2537) or mevalonate (Sigma #M4667) at the concentrations indicated in Fig. 2E,F. Images were taken daily and half-media changes performed every second day, with half-concentrations of the treatments used when refreshing media.

### QUANTIFICATION AND STATISTICAL ANALYSIS

#### Quant-Seq analysis

Fastq files were uploaded to and processed through the cloud-based BlueBee Genomics Platform (https://www.bluebee.com/quantseq). Briefly, raw reads were trimmed with Bbduk, aligned with STAR Aligner, and counted with HTSeq-count to yield gene counts. Differential expression analyses were performed with the DESeq2 pipeline, which compared counts from each set of duplicates at different timepoints to counts from the day 0 timepoints. Additional, custom analysis pipelines were devised in R. We developed a simple web application with the Shiny R package that enables users to visualize normalized read counts and expression fold change (relative to day 0) throughout neuronal differentiation for a queried gene. The web application can be accessed via kampmannlab.ucsf.edu/ineuron-rna-seq.

#### Primary screen analysis

We developed a bioinformatics pipeline, MAGeCK-iNC (MAGeCK including Negative Controls) for large-scale functional genomics analysis, which we made publicly available (kampmannlab.ucsf.edu/mageck-inc). First, raw sequencing reads from next-generation sequencing were cropped and aligned to the reference using Bowtie (Langmead et al., 2009) to determine sgRNA counts in each sample. Next, counts files of two samples subject to comparison were input into MAGeCK and log2 fold changes (LFCs) and P values were calculated for each sgRNA using the ‘mageck test –k’ command. Following that, gene level knockdown phenotype scores were determined by averaging LFCs of the top 3 sgRNAs targeting this gene with the most significant P values. The statistical significance for each gene was determined by comparing the set of P values for sgRNAs targeting it with the set of P values for non-targeting control sgRNAs using the Mann-Whitney U test, as described previously (Kampmann et al., 2013, 2014). To correct for multiple hypothesis testing, we first performed random sampling of 5 with replacement from non-targeting control sgRNAs to generate ‘negative-control-quasi-genes’ and calculated knockdown phenotype scores and P values for each of them. Then, we calculated the hit strength, defined as the product of knockdown phenotype score and –log (p value), for all genes in the library and for ‘negative-control-quasi-genes’ generated above. Based on the distribution of all the products, a cutoff value was chosen to make sure the false-discovery rate (FDR) is less than 0.05. To find enriched annotations within hit genes, Gene Set Enrichment Analysis (GSEA) was performed for Day 10 iPSCs and Day 28 neurons using the fgsea package in R (Sergushichev, 2016).

#### Pooled validation screen analysis

sgRNA counts for each sample were determined as in primary screen. Subsequently, knockdown phenotype scores for each sgRNA were calculated as LFCs of sgRNA counts between two samples and were normalized by subtracting the median of non-targeting control sgRNAs. LFCs were averaged for samples with replicates. Gene-level knockdown phenotype score was determined as the mean of knockdown phenotype scores of all sgRNAs targeting this gene.

#### CROP-Seq analysis

Cell Ranger (version 2.2.0, 10X Genomics) with default parameters was used to align reads and generate digital expression matrices from single-cell sequencing data. To map sgRNA transcripts together with other mRNA transcripts to individual cells, a custom reference was generated by extending the human genome assembly (Ensembl GRCh38 release) with ‘quasi-genes’ representing sgRNA-containing transcripts (one sgRNA sequence per quasi-gene with 250bp upstream and 230bp downstream sequences). Sequencing results of sgRNA-enrichment libraries were analyzed using methods previously described (Hill et al., 2018) to further facilitate sgRNA identity assignment.

For a given cell, sgRNA(s) whose UMI counts were greater than 4 standard deviations of the mean UMI counts of all sgRNAs were assigned to that cell as its identity. Cells with only one assigned sgRNA were retained for further analysis. The Scater package (McCarthy et al., 2017) implemented in R was used to analyze the digital expression matrices including normalization, quality control and filtering.

The mean reads per cell was around 84,000 for iPSCs and 91,000 for neurons. Median number of genes detected per cell was around 5,000 for iPSCs and 4,600 for neurons. After quality control, a single sgRNA could be assigned to ~15,000 iPSCs and ~8,400 neurons

For each target gene, the top 50% cells with best on-target knockdown were retained. Differential gene expression analysis was performed between each gene knockdown group (cells assigned by targeting sgRNAs of that gene) and control group (cells assigned by non-targeting control sgRNAs) using the R package edgeR (Robinson et al., 2010) treating each cell as one replicate.

For Fig. 4C, relative expression of each gene was calculated as z-normalized expression with respect to the mean and standard deviation of that gene in the control group:

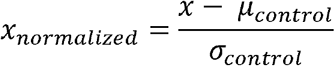

The top 20 most significantly altered genes were selected for each gene knockdown group and merged together to form the signature gene list. Gene knockdown groups were hierarchically clustered based on their relative expression of the signature genes, using Cluster 3.0 (Eisen et al., 1998) and visualized using Java TreeView (Saldanha, 2004).

For Fig. 6E, GSEA was performed using the web tool WebGestalt (Zhang et al., 2005). For Fig. S5C,D, the similarity score of transcriptome changes between two gene knockdown groups, A and B, was calculated as follows:

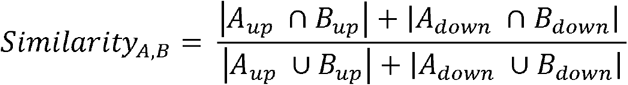

*A_up_* and *B_up_* denote for the significantly upregulated genes (p_adj_<0.01) in A and B, while *A_down_* and *B_down_* denote for the significantly downregulated genes (p_adj_<0.01) in A and B.

#### Longitudinal imaging data analysis

A CellProfiler (Carpenter et al., 2006) pipeline was developed to analyze longitudinal imaging data. For iPSC growth and i^3^Neuron survival experiments, sgRNA+ cells were recognized as nuclear-green+ blue+ objects and the total number of sgRNA+ cells was quantified for every image. iPSC growth and i^3^Neuron survival were calculated as the ratio of sgRNA+ cell number at different time points to that of day 1 of imaging. For neurite morphology analysis, neurites of sgRNA+ cells were first enhanced by the EnhanceOrSuppressFeatures and EnhanceEdges modules, and then skeletonized by the Morph module. Following that, MeasureObjectSkeleton module was implemented to measure neurite length, number of branches and number of trunks for individual neurons. The mean values of the above measurements of all sgRNA+ neurons were calculated for each image.

To integrate all image analysis data, we generated a panel of imaging phenotypes for a given sgRNA, including neurite length, number of neurite branches, number of neurite trunks, neuronal survival and iPSC growth at different time points. For Fig. 7E, the percentage changes of imaging phenotypes compared to the mean of non-targeting control sgRNAs were calculated for each sgRNA. Most of genes in the imaging experiment were targeted by two sgRNAs (some genes missed one sgRNA during experiment process), and a gene was discarded if it was targeted by two sgRNAs and the correlation of the two sgRNAs was less than 0.8. All remaining sgRNAs were hierarchically clustered based on the Pearson correlation of their percentage changes of imaging phenotypes, using Cluster 3.0 (Eisen et al., 1998) and visualized using Java TreeView (Saldanha, 2004).

#### Calcium Imaging Analysis

Representative GCaMP6m movies for each cell group were chosen manually, after viewing all movies for each group. Representative movies before and after CNQX addition were merged sequentially and aligned using Nikon Elements software. For each merged and aligned movie, ROIs were drawn manually with Fiji (Schindelin et al., 2012) around every clearly visible cell body, and the mean gray value was measured for each ROI in each frame. For each ROI, ΔF/F was calculated using the average of the 5 frames with the lowest values as the baseline.

### DATA AND SOFTWARE AVAILABILITY

RNA sequencing data sets generated in this study are available on NCBI GEO as dataset GSE124703 (https://www.ncbi.nlm.nih.gov/geo/query/acc.cgi?acc=GSE124703).

Expression levels of genes of interest at different time points during neuronal differentiation can be visualized interactively at kampmannlab.ucsf.edu/ineuron-rna-seq.

Longitudinal imaging data files will be made available on request to the lead author. The MAGeCK-iNC bioinformatics pipeline is available at kampmannlab.ucsf.edu/mageck-inc.

The CellProfiler pipeline for analysis of neuronal longitudinal imaging data will be made available on request to the lead author, and will also be submitted to the CellProfiler depository of published pipelines (https://cellprofiler.org/examples/published_pipelines.html) upon publication.

## SUPPLEMENTAL INFORMATION TITLES AND LEGENDS

**Table S1. Phenotypes from the primary screen** (related to Fig. 2)

Phenotypes from the primary screens (Day 10 iPSCs, Day 14 neurons, Day 21 neurons, Day 28 neurons) are listed for all genes targeted H1 library (see Methods for details). Columns are: Targeted transcription start site, targeted gene, Knockdown phenotype, P value, and the product of phenotype x –log10(P value).

**Table S2. Pooled validation screen sgRNAs and phenotypes** (related to Fig. 3)

Protospacer sequences and phenotypes from the validation screens are listed for all sgRNAs from the pooled validation library (see Methods for details). Columns are: sgRNA short name (as used throughout the study), sgRNA long names (from the original H1 library), protospacer sequences, and phenotypes from the different screens in the following columns.

**Table S3. Differentially expressed genes from the CROP-Seq screen** (related to Figs. 4–6)

The first two tabs provide the numerical values underlying the heatmaps in Fig. 4C. Columns: genes targeted by CRISPRi. Rows: differentially expressed genes. The following tabs list changes in gene expression for *MAP3K12, UQCRQ, UBA1* or *MAT2A* knockdown versus non-targeting sgRNAs. See Methods for details.

**Movie S1. Calcium imaging of i^3^N neurons on Day 28** (related to Fig. 1)

2 time lapse movies acquired at approximately 5 fps are concatenated, before and 1 min after addition of 50 uM CNQX.

**Movie S2. Calcium imaging of CRISPRi i^3^N neurons transduced with a non-targeting sgRNA on Day 28** (related to Fig. 1)

2 time lapse movies acquired at approximately 5 fps are concatenated, before and 1 min after addition of 50 uM CNQX.

**Movie S3. Time-lapse of longitudinal imaging for iPSC-derived neurons expressing a non-targeting control sgRNA** (related to Fig. 6)

The ten frames represent Days 1, 2, 3, 4, 5, 6, 8, 10, 13, and 16 of imaging. Red: cytosolic mScarlet. Blue: nuclear-localized BFP marker from sgRNA construct. The movie is shown at a speed of 4 fps.

**Movie S4. Time-lapse of longitudinal imaging for iPSC-derived neurons expressing a sgRNA targeting *UQCRQ*** (related to Fig. 6)

The ten frames represent Days 1, 2, 3, 4, 5, 6, 8, 10, 13, and 16 of imaging. Red: cytosolic mScarlet. Blue: nuclear-localized BFP marker from sgRNA construct. The movie is shown at a speed of 4 fps.

## Notes

#### Summary of Updates

Additional control experiments were conducted to demonstrate that CRISPRi machinery and/or sgRNAs did not cause DNA damage and did not affect neuronal differentiation, survival, axon growth, and neuronal activity as monitored by calcium imaging. Additional bioinformatics analyses and Figures were included to provide additional insights.

