## Supplemental Figures and Table S4 for "CRISPR interference-based platform for multimodal genetic screens in human iPSC-derived neurons"

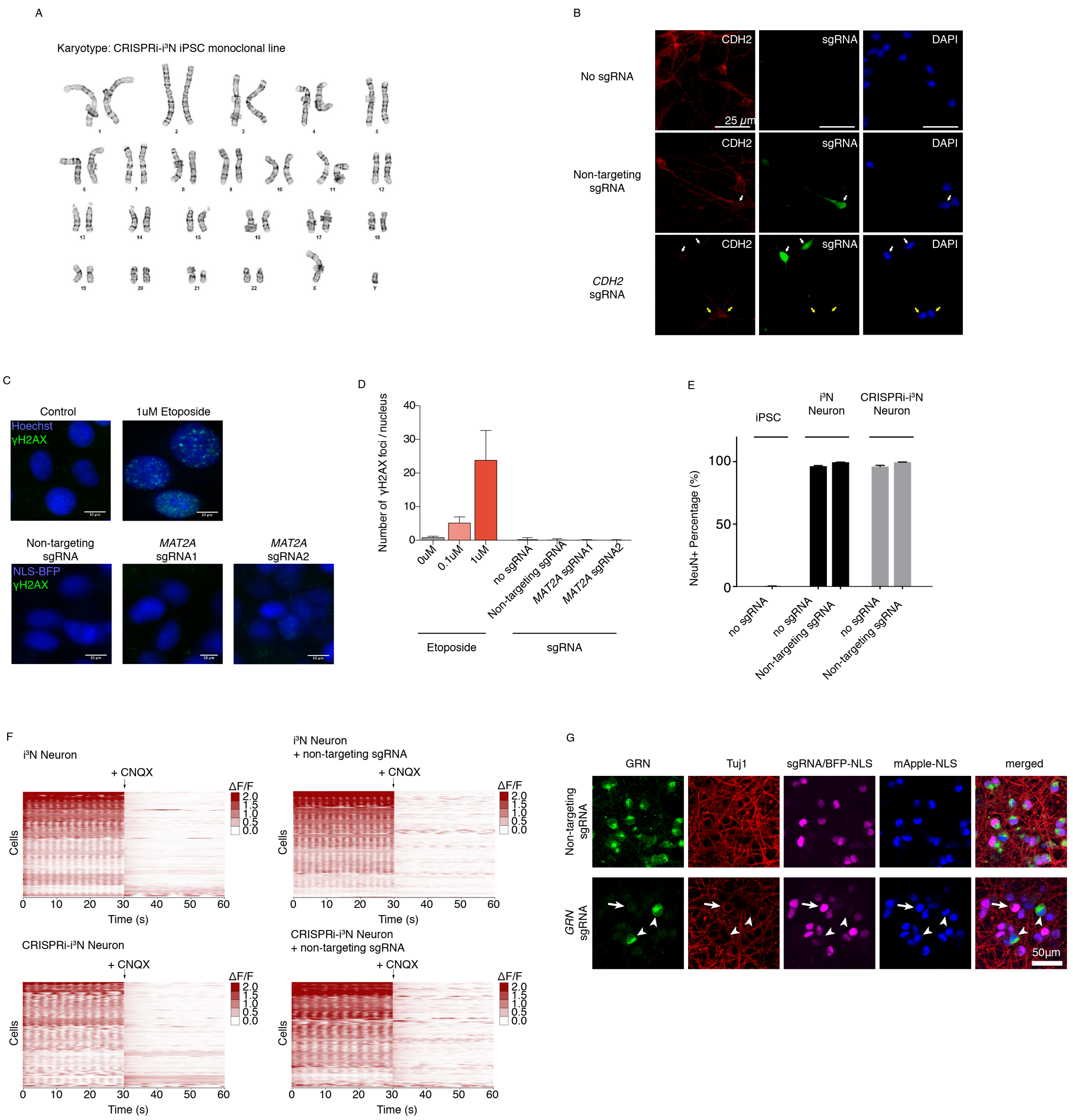


**Figure S1. Normal karyotype, neuronal differentiation and activity and CRISPRi activity of the CRISPRi- i$^3$N iPSC monoclonal line** (related to Fig. 1)

**(A)** Karyotyping of the monoclonal CRISPRi- i$^3$N iPSC line confirmed a normal male karyotype.

**(B)** Knockdown of N-cadherin (*CDH2*) in iPSC-derived neurons on Day 18 monitored on the protein level by immunofluorescence (IF) microscopy. White arrows mark cells infected with a lentiviral plasmid expressing an sgRNA and GFP (green). *Top row*, uninfected cells. *Middle* *row*, non-targeting negative control sgRNA. N-cadherin signal (IF, red) is similar in infected and uninfected cells. *Bottom row*, sgRNA targeting N-cadherin (*CDH2*). For the infected cells (white arrows), N-cadherin signal (IF, red) is substantially reduced compared with neighboring uninfected cells (yellow arrows). Nuclear counterstain DAPI is shown in blue.

(**C**,**D**) DNA damage detected by γH2AX staining. CRISPRi-i$^3$N iPSCs treated with vehicle or 1μM etoposide for 6 hours or infected with non-targeting sgRNA or two different *MAT2A* sgRNAs for 48 hours were fixed and immunostained using an γH2AX antibody. (C) Representative images of γH2AX staining (Green). γH2AX foci can be observed in cells treated with etoposide, but not in cells with no treatment or with sgRNAs. Nuclei are visualized by Hoechst stain (*top row*, control and etoposide treatment) or NLS-BFP (*bottom row*, sgRNAs). (D) Quantification of the number of γH2AX foci per nucleus using CellProfiler. Mean and standard deviations for 6 replicates wells are shown.

(**E**) Quantification of the percentage of NeuN positive cells in i$^3$N iPSCs, i$^3$N neurons and CRISPRi-i$^3$N neurons with or without transduction of a non-targeting sgRNA. Mean and standard deviations for replicates are shown.

(**F**) Heatmaps showing neuronal activity by GCaMP6m calcium imaging in i$^3$N neurons and CRISPRi-i$^3$N neurons with or without a non-targeting sgRNA.

(**G**) Knockdown of progranulin (*GRN*) in CRISPRi-NCRM5 neurons on Day 5 detected at the protein level by immunostanining. *Top row*, non-targeting control sgRNA. *Bottom row*, sgRNA targeting progranulin. For cells infected with progranulin-targeting sgRNA (indicated by arrow), progranulin signal (IF, green) is substantially reduced compared with cells infected with non-targeting sgRNA or cells not infected with sgRNA (indicated by arrowheads). The neuronal marker Tuj1 (IF, red), an sgRNA marker (BFP-NLS, purple) and a nuclear marker (mApple-NLS, blue) are shown.

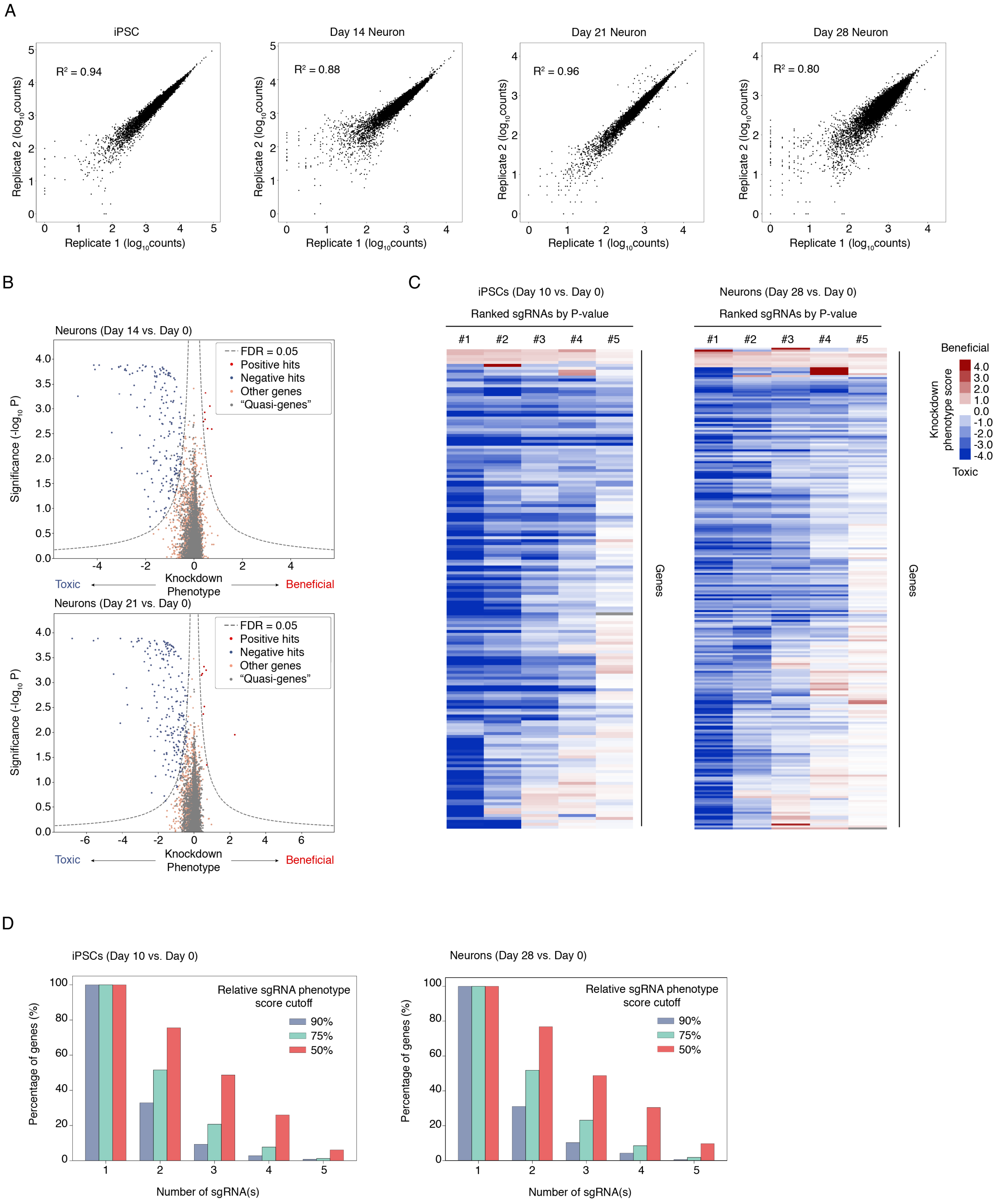


**Figure S2. Characterization of results from massively parallel screens for essential genes in iPSCs and neurons** (related to Fig. 2)

**(A)** Correlation of read counts from next-generation sequencing for individual sgRNAs between experimental replicates.

**(B)** Volcano plots summarizing knockdown phenotypes and statistical significance (Mann-Whitney U test) for genes targeted in the pooled screen in iPSC-derived neurons. *Top*, phenotypes for survival between Day 0 and Day 14. *Bottom*, phenotypes for survival between Day 0 and Day 21. Dashed lines represent the cutoff for hit genes, which was defined based on the product of phenotype and $-\log_{10}$(P value) at an empirically determined false discovery rate of 0.05 (see Methods).

**(C)** Heatmaps showing knockdown phenotype scores of all 5 sgRNAs (x-axis) for every hit gene (y-axis) in iPSCs (*left*) and iPSC-derived neurons (*right*). The five sgRNAs for a given gene are ranked by their p-values and are shown from left to right.

**(D)** Bar graphs summarizing the percentage of hit genes that have a certain number of sgRNAs (x-axis) showing a knockdown phenotype score above a given activity cutoff in iPSCs (left) and iPSC-drived neurons (right). For each sgRNA targeting a given gene, a relative sgRNA phenotype score was calculated by dividing its knockdown phenotype score by that of the most significant sgRNA targeting the same gene. Cutoffs are set based on the relative sgRNA phenotype scores.

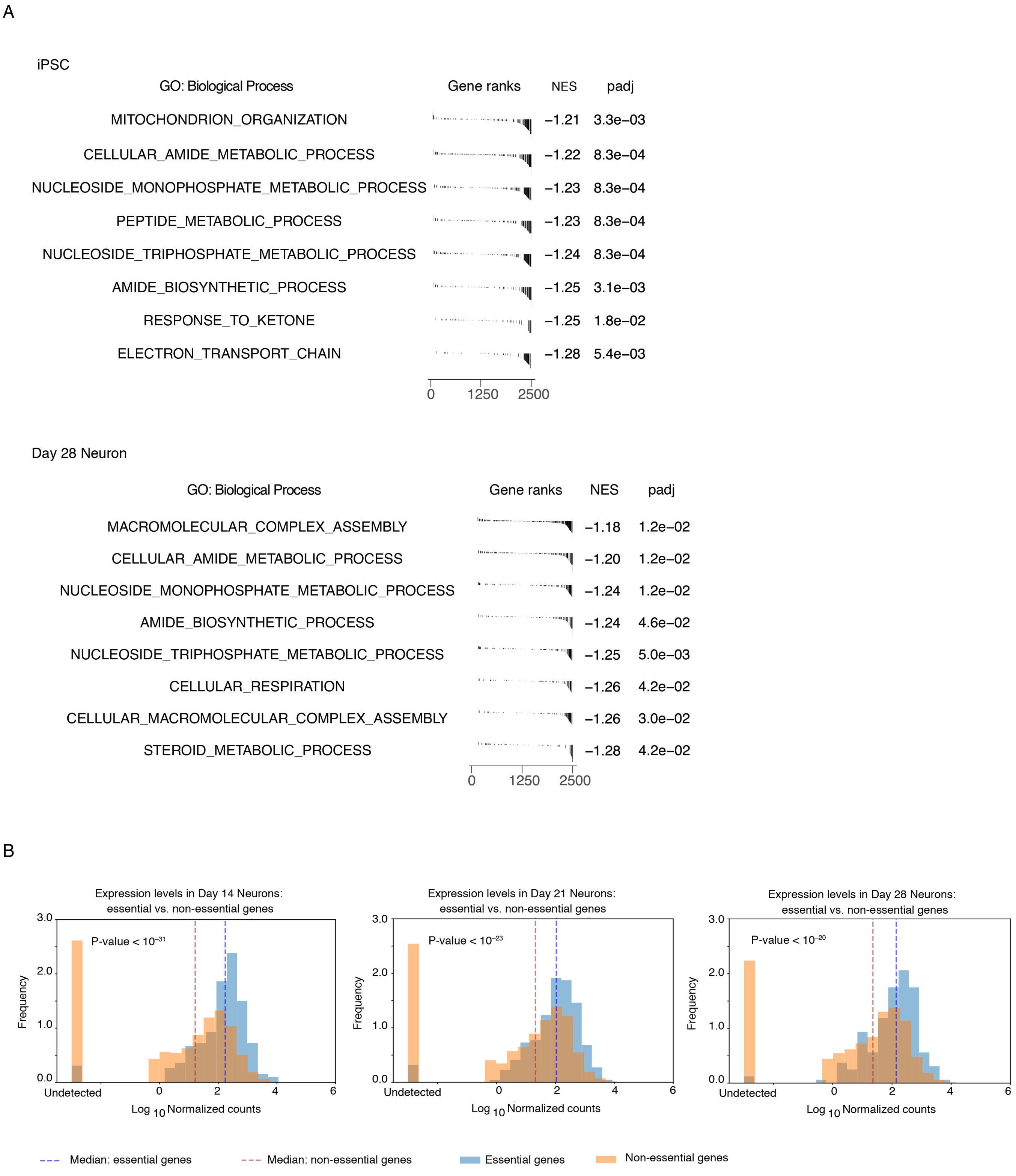


**Figure S3. Functional analysis of hit genes from primary survival screens in iPSCs and iPSC-derived neurons** (related to Fig. 2)

(**A**) Gene set enrichment analysis (GSEA) for hit genes from the screens. Significantly enriched GO terms for biological processes (BH-adjusted P value<0.05: 100,000 permutations) are shown for essential genes in iPSCs (Day 0 vs. Day 10) and neurons (Day 0 vs. Day 28). NES, normalized enrichment score.

(**B**) Expression levels of essential and non-essential genes in neurons at Day 14, 21, and 28 of differentiation were plotted as the distributions of $\log_{10}$ normalized counts from Quant-Seq. Median levels of expression were indicated. P values were calculated using one-sided Mann-Whitney U test.

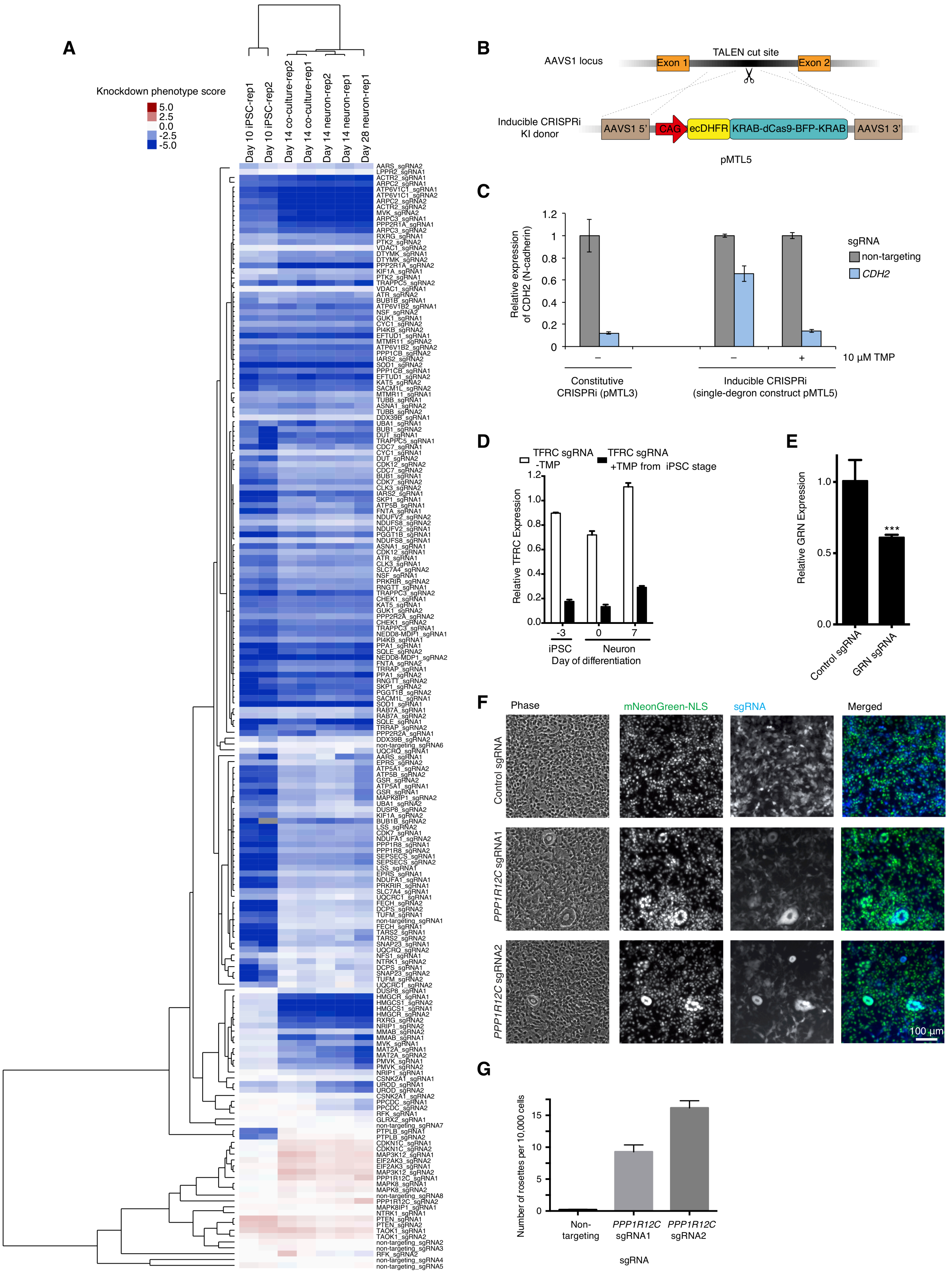


**Figure S4. Pooled validation of hit genes from the primary screen** (related to Fig. 3)

(**A**) Heatmap showing relative knockdown phenotype scores of sgRNAs (rows) from the validation screens (columns, both experimental replicates, expect for Day 28 neurons, for which the second replicate sample was accidentally lost during sample preparation). Both sgRNAs and screens were hierarchically clustered based on Pearson correlation.

(**B**) Construct pMTL5 for the expression of CRISPRi machinery (dCas9-BFP-KRAB) fused to an N-terminal DHFR degron from the AAVS1 safe-harbor locus.

(**C**) Characterization of inducible CRISPRi activity in iPSCs with integrated single-degron CRISPRi machinery (construct pMTL5), compared to an equivalent constitutive construct lacking the degron (pMTL3). iPSCs were transduced with a lentiviral sgRNA expression construct containing a non-targeting sgRNA (grey bars) or an sgRNA targeting *CDH2* (encoding N-cadherin, blue bars). Cells were cultured in the presence or absence of 10 µM trimethoprim (TMP), and cell-surface levels of N-cadherin were quantified by immunofluorescence flow cytometry 48 hours after transduction. Expression levels relative to non-targeting sgRNA are shown. Mean of two experimental replicates, error bars indicate standard deviation.

**(D,E)** Characterization of inducible CRISPRi activity in iPSCs and neurons with integrated double-degron CRISPRi machinery (construct pRT029). (D) Knockdown of the transferrin receptor (*TFRC*) in iPSCs and iPSC-derived neurons in the presence or absence of trimethoprim (TMP). iPSCs expressing the inducible CRISPRi machinery were lentivirally infected with an sgRNA targeting *TFRC* or a non-targeting negative control sgRNA. Infected iPSCs were cultured and differentiated into neurons in the presence of 20 µM TMP from iPSC stage or in the absence of TMP. Cells from both conditions were harvested at different days and levels of *TFRC* as well as *ACTB* mRNAs were quantified by qPCR. After normalizing each sample by *ACTB* mRNA levels, ratios of *TFRC* mRNA were calculated for cells expressing the *TFRC*-targeting sgRNA versus cells with the non-targeting sgRNA cultured in the same condition. Mean and standard deviations for replicates are shown. (E) qPCR result showing the conditional knockdown of *GRN* in iPSC-derived neurons. iPSCs expressing the dual-degron inducible CRISPRi machinery were lentivirally infected with an sgRNA targeting *GRN* or a non-targeting negative control sgRNA. Infected iPSCs were differentiated into neurons. The neurons were cultured in neuronal medium with the presence or absence of 20 µM TMP. Day 7 neurons from both conditions were harvested and levels of *GRN* as well as *ACTB* mRNAs were quantified by qPCR. After normalizing each sample by *ACTB* mRNA levels, ratios of *GRN* mRNA were calculated for cells expressing the *GRN*-targeting sgRNA versus cells with the control sgRNA cultured in the same condition. Mean and standard deviations for replicates are shown. One-tailed t-test was applied (*** $P<0.001$).

(**F,G**) Inhibition of the neuronal differentiation of CRISPRi-i$^3$N iPSCs by sgRNAs targeting *PPP1R12C*. (F) Phase contrast (first column) and fluorescence (second and third column) micrographs of partially differentiated neuron populations infected with an expression construct for sgRNAs (non-targeting control sgRNA, top row, or two different sgRNAs targeting *PPP1R12C,* middle and bottom row) and a BFP marker. Nuclei are visualized by expression of mNeonGreen-NLS (second column, green in merged images). Cells expressing sgRNAs are marked by cytosolic BFP (third column, blue in merged images). Images were acquired following six days of iPSC proliferation post-infection and three days of doxycycline-induced differentiation. (G) Quantification of undifferentiated colonies in a repeat experiment. Rosettes were counted manually by an individual blinded to the experimental conditions. The total number of cells was counted by Nikon Elements software bright spot detection module for green nuclei. Mean and SEM for replicate wells are shown (n = 6). One-way ANOVA and multiple comparisons were applied (****P<0.0001).

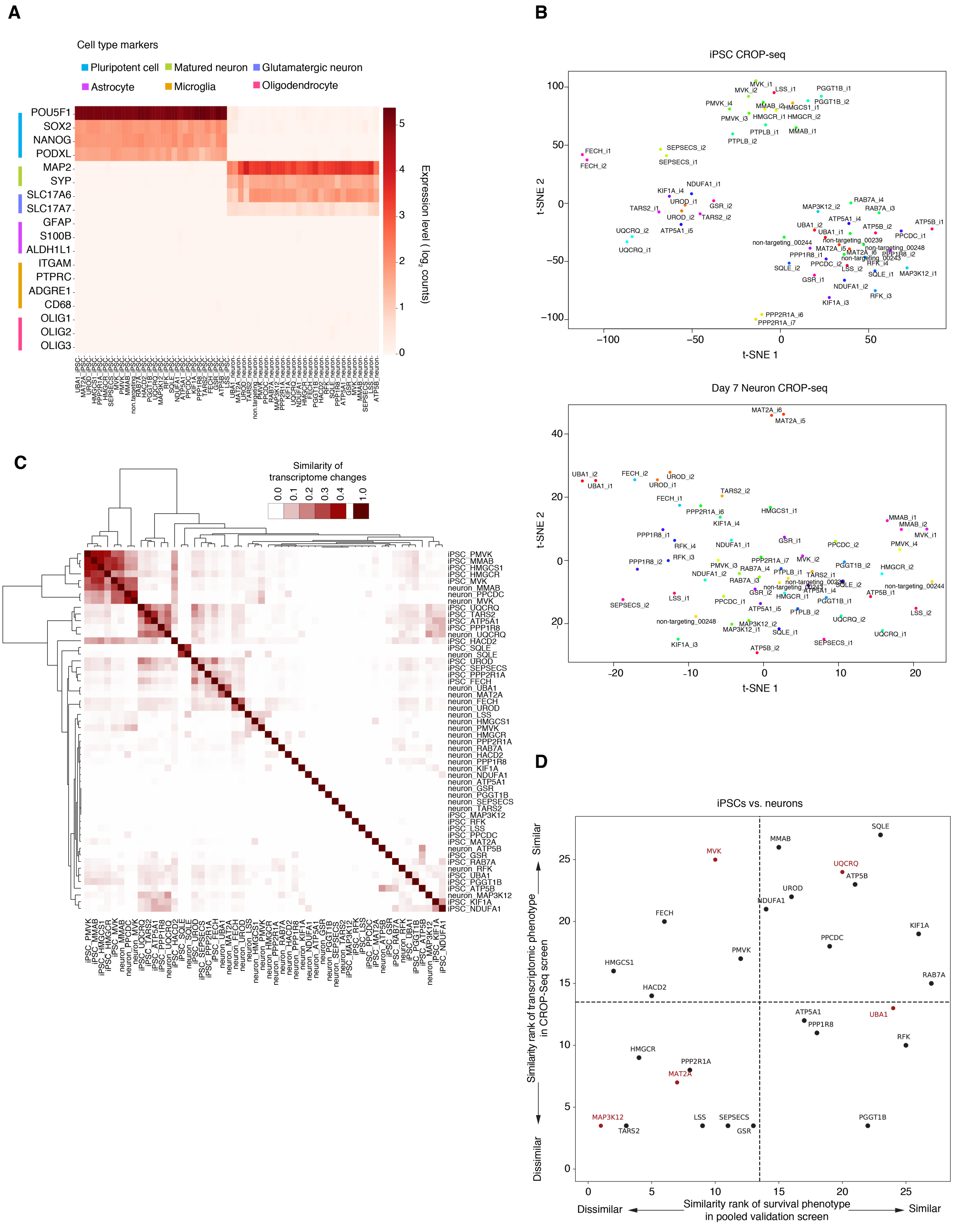


**Figure S5. Characterization of CROP-Seq screen results** (related to Fig. 4)

(**A**) Heatmap showing the expression levels of marker genes for different cell types in different gene knockdown groups in iPSCs and Day 7 neurons obtained from the CROP-seq screen. The shade of red indicates the mean expression level of a given gene for all cells in that gene knockdown group.

(**B**) Transcriptomes of different sgRNA groups in iPSCs (left) and neurons (right) were visualized with t-Distributed stochastic neighbor embedding (t-SNE), colored by target genes.

(**C**) Pairwise similarities of transcriptome changes of different gene knockdown groups across iPSCs and neurons were determined based on the numbers of overlapping and total transcripts that were significantly altered in two groups (see Methods for details). Gene knockdown groups were hierarchically clustered based on Pearson correlation. The shade of red indicates the similarity.

(**D**) Comparison of differences between iPSCs and neurons in knockdown phenotypes for survival and transcriptomic responses. Genes were ranked by similarities between iPSCs and neurons in their knockdown phenotypes in terms of survival (x-axis) or transcriptomic response (y-axis). Dashed lines indicate the middle rank positions. Genes selected for further discussion are labeled in red.

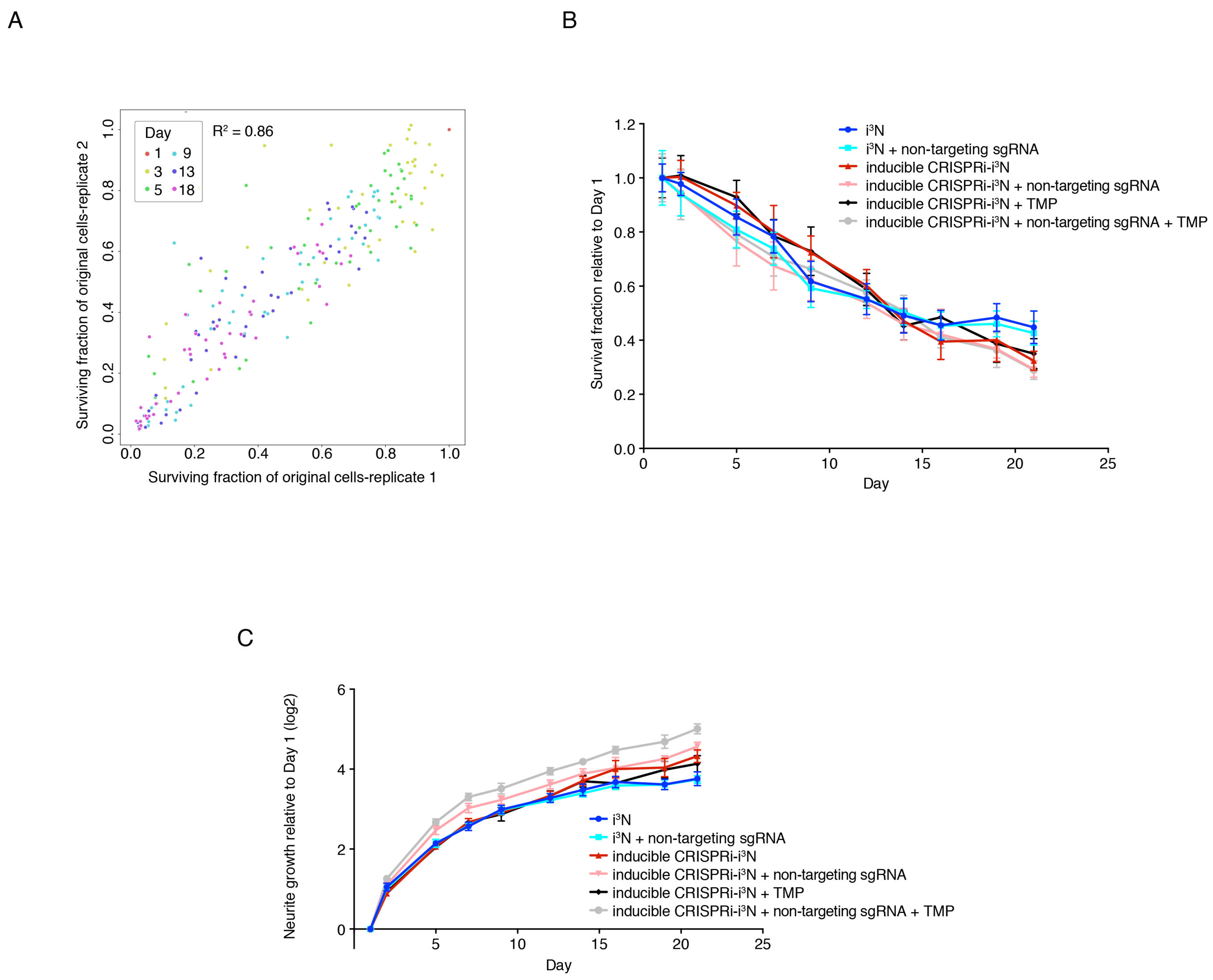


**Supplemental Fig. S6. Reproducibility of longitudinal imaging results** (related to Fig. 7)

(**A**) Scatter plot showing surviving fractions of neurons with different sgRNAs at different days of differentiation (shown in different colors) relative to Day 1 in two independent sets of imaging experiments. Each dot represents one sgRNA in a certain time point. Coefficient of determination ($R^2$) is indicated.

(**B,C**) Survival (B) and neurite growth (C) of $i^3N$ neurons and inducible CRISPRi $i^3N$ neurons with or without non-targeting sgRNA in the absence or presence of TMP, quantified from longitudinal imaging data. Mean and standard deviations for 6 replicate wells are shown.

| Name | Sequence | Sequence (5' to 3') | Use |
|---|---|---|---|
| B-actin For | entire | ACCTTCTACAATGAGCTGCG | qPCR |
| B-actin Rev | entire | CCTGGATAGCAACGTACATGG | qPCR |
| GAPDH For | entire | ATGCCTCCTGCACCACCAAC | qPCR |
| GAPDH Rev | entire | GGGGCCATCCACAGTCTTCT | qPCR |
| CDH2 For | entire | CCCAAGACAAAGAGACCCAG | qPCR |
| CDH2 Rev | entire | GCCACTGTGCTTACTGAATTG | qPCR |
| TFRC For | entire | ACTTGCCCAGATGTTCTCAG | qPCR |
| TFRC Rev | entire | GTATCCCTCTAGCCATTCAGTG | qPCR |
| GRN For | entire | TCCAGAGTAAGTGCCTCTCCA | qPCR |
| GRN Rev | entire | TCACCTCCATGTCACATTTCA | qPCR |
| Enrichment_ PCR_1_For | entire | TCGCACGGACTTGTGGGAGAAG | sgRNA enrichment PCR |
| Enrichment_ PCR_1_Rev | entire | CTACACGACGCTCTTCCGATCT | sgRNA enrichment PCR |
| Enrichment_ PCR_2_For | entire | GTGACTGGAGTTCAGACGTGTGCTCTTCC GATCTAGTATCCCTTGGAGAACCACCTTG | sgRNA enrichment PCR |
| Enrichment_ PCR_2_Rev | entire | AATGATACGGCGACCACCGAGATCTACAC TCTTTCCCTACACGACGCTC | sgRNA enrichment PCR |
| Enrichment_ PCR_3_For | entire | CAAGCAGAAGACGGCATACGAGAT-[index]-GTGACTGGAGTTCAG | sgRNA enrichment PCR |
| CLYBL _sgRNA | protospacer | ATGTTGGAAGGATGAGGAAA | sgRNA targeting the human CLYBL intragenic safe harbor locus |
| non-targeting control sgRNA_1872 | protospacer | GTCCACCCTTATCTAGGCTA | Figs. 1C,D,E,G, S1B, S4C,D,E |
| non-targeting control sgRNA_1873 | protospacer | GGACTAAGCGCAAGCACCTA | Fig. 1E,F,G, S4F,G |
| non-targeting control sgRNA-3180 | protospacer | GAGACGAGGACATGTGTAGC | Figs. S1C,D,E, 2E, S6B |
| non-targeting control sgRNA-3181 | protospacer | GTACCACCCAAACGATAACG | Fig. S1C,D |
| CDH2_sgRNA | protospacer | GGGGCCGAGCGAAGAGCCGG | Figs. S1B, S4C |
| TFRC_sgRNA | protospacer | GCTCAGAGCGTCGGGATATC | Figs. 1C, S4D |
| GRN_sgRNA | protospacer | GTGCCCAAGGACCGCGGAGT | Figs. 1F,G, S1G, S4E |
| UBQLN2_ sgRNA | protospacer | GTGCGCGCGGTCGGATCACA | Fig. 1D,E |
| HMGCR_ sgRNA_3018 | protospacer | GCACCTCCAGATCTCACTAG | Fig. 2F |
| MAT2A_sgRNA_1 | protospacer | GTCTACTCGTAGCAGGCGGG | Fig. S1C,D |
| MAT2A_sgRNA_2 | protospacer | GTGGCCTTACGAAGGAGCAG | Fig. S1C,D |
| PPP1R12C_sgRNA_1 | protospacer | GCCCAGGGCCAGGAACGACG | Fig. S4F,G |
| PPP1R12C_sgRNA_2 | protospacer | GGCACAGGCCGCCAGGAACT | Fig. S4F,G |

**Table S4. Oligonucleotide sequences for primers and sgRNAs** (related to STAR Methods)
